# Male and female bees show large differences in floral preference

**DOI:** 10.1101/432518

**Authors:** Michael Roswell, Jonathan Dushoff, Rachael Winfree

**Affiliations:** Graduate program in ecology and evolution, Rutgers University, 14 College Farm Road, New Brunswick, NJ 08904; Department of biology, McMaster University, 1280 Main St. West, Hamilton, Ontario ON L8S 4K1; Department of ecology, evolution, and natural resources, Rutgers University, 14 College Farm Road, New Brunswick, NJ 08904

**Keywords:** dimorphism, dissimilarity, Morisita-Horn, phenology, plant-pollinator interaction, pollination, pollinator habitat, preference

## Abstract

1. Intraspecific variation in foraging niche can drive food web dynamics and ecosystem processes. Field studies and theoretical analysis of plant-pollinator interaction networks typically focus on the partitioning of the floral community between pollinator species, with little attention paid to intraspecific variation among plants or foraging bees. In other systems, male and female animals exhibit different, cascading, impacts on interaction partners. Although the foraging ecology of male bees is little known, we expect foraging preferences to differ between male and female bees, which could strongly impact plant-pollinator interaction outcomes.
2. We designed an observational study to evaluate the strength and prevalence of sexually dimorphic foraging preferences in bees.
3. We observed bees visiting flowers in semi-natural meadows in New Jersey, USA. To detect differences in flower use against a shared background resource availability, we maximized the number of interactions observed within narrow spatio-temporal windows. To distinguish observed differences in bee use of flower species, which can reflect abundance patterns and sampling effects, from underlying differences in bee preferences, we analyzed our data with both a permutation-based null model and random effects models.
4. We found that the diets of male and female bees of the same species were often as dissimilar as the diets of bees of different species. Furthermore, we demonstrate differences in preference between male and female bees, and identify plant species that are particularly attractive to each sex. We show that intraspecific differences in preference can be robustly identified within interactions between hundreds of species, without precisely quantifying resource availability, and despite high phenological turnover.
5. Given the large differences in flower use and preference between male and female bees, ecological sex differences should be integrated into studies of bee demography, plant pollination, and coevolutionary relationships between flowers and insects.

## Introduction

Intraspecific variation in traits and behavior, including foraging niche, has important consequences for species interactions and conservation (Bolnick et al., 2011; Durell, 2000). Sexual dimorphism is a large source of individual niche variation, and an important factor in plant-animal interactions, such as seed dispersal (Zwolak, 2018). Sexual dimorphism underlies adaptation, speciation, and the way in which animals exploit their ecological niche (Butler, Sawyer, & Losos, 2007; Temeles, Miller, & Rifkin, 2010). Morphological, behavioral, and life-history dimorphisms can also drive the form and function of ecosystems, for example when predator sex ratio drives the community composition of lower trophic levels, shown experimentally (Start & De Lisle, 2018) and in nature, where responses extended to water chemistry as well (Fryxell, Arnett, Apgar, Kinnison, & Palkovacs, 2015).

Though ecological dimorphisms were first studied in vertebrates (Selander, 1966), they are common across taxa, including insects (Shine, 1989). Surprisingly, in bees (Hymenoptera, Apoidea) for which both foraging (P. Willmer, 2011) and sexual dimorphism (Alcock et al., 1978) have been well studied, sexually dimorphic foraging has rarely been documented. Intraspecific variation in floral preference is well known in social (Heinrich, 1979) and to a lesser extent, solitary bee species (Bruninga-Socolar, Crone, & Winfree, 2016; Tur, Vigalondo, Trøjelsgaard, Olesen, & Traveset, 2014), yet most community-level studies focus on species-level interaction networks, and furthermore, on the foraging preferences of only female bees.

Male bees differ from their better-studied female counterparts in their life history and ecology. Female bees construct, maintain, provision, and defend nests, whereas male bees primarily seek mates (P. G. Willmer & Stone, 2004). Both sexes drink floral nectar for their own caloric needs, but only females collect pollen to provision young, and thus forage at greater rates. While the pollen from each flower species (the term we use throughout for the flowers from a species of plant) tends to be morphologically and nutritionally distinct, interspecific variation in the chemical composition of nectar is comparatively subtle (Cane & Sipes, 2006). Thus, we expect the foraging ecology, including floral preferences, of male and female bees to differ as well.

Male bees prove to be important pollinators when studied, both in specialized oil- or scent-collecting pollination systems (Eltz et al., 2007; Etl, Franschitz, Aguiar, Schönenberger, & Dötted, 2017; Janzen, 1971) and also when males are foraging for nectar and pollen (Cane, 2002; Cane, Sampson, & Miller, 2011; Ogilvie & Thomson, 2015). Male bees may also be particularly relevant for bee conservation. Males may be limiting in declining populations, either because genetic diversity is necessary for the development of female offspring as a result of complementary sex determination, or because mate or sperm limitation results from poor male condition (Elias, Dorn, & Mazzi, 2010; Straub et al., 2016). As the dispersing sex in most bee species, males may be crucial for gene flow and metapopulation persistence even when they are not locally limiting (López-Uribe, Morreale, Santiago, & Danforth, 2015; Ulrich, Perrin, & Chapuisat, 2009).

Foraging niche is only partly described by resource use. Indeed, resource preferences may be more important than use alone in many contexts, including conservation. Preference—the use of a resource in excess of its relative availability—is challenging to measure, because both resource use and availability must be known. Floral resource availability for pollinators is particularly hard to quantify outside an experimental context because the appropriate scale and units of floral resource availability are unclear. The composition, amount, and supply rate of pollen and nectar per flower, the number of flowers per inflorescence, of inflorescences per individual, and the number and distribution of individual plants over the square kilometers of a bee’s foraging range are all important components of availability (Hicks et al., 2016). Furthermore, floral availability can change rapidly over time. However, differences in flower use between bees foraging at the same place and time indicate differences in preference, which may occur between species, or between individuals of the same species.

In this study, we assess differences between floral preferences of male and female bees in the wild. We collected bees foraging on flowers in meadows in New Jersey, USA. In order to observe preference differences, we collected as many individuals as possible during replicated, short (3-day) windows, during which we assumed floral availability and bee abundance were constant at each site. We compare the species composition of flowers visited by males and females of the most common bee species across the entire study as a naïve measure of differences in preference between the sexes. Then, using random effects models, we assess when differential flower species *use* by male and female bees likely arises from sex-specific floral *preference*, as opposed to shifting overlap between foragers and floral resources (i.e. changes in *availability* without differences in preference). Specifically, we ask

1. How much do male and female bee diets overlap?
2. To what degree are particular flower species disproportionately visited by bees of one sex?
3. To what extent are differences in floral use driven by preference, rather than phenological differences between male and female bees?

## Materials and Methods

### Study design and data collection

Because absolute preference is nearly impossible to observe outside of an experiment, we designed our study to reveal differences in preference between groups of bees. In order to collect a large number of males and females from many native bee species, we selected six meadows (sites) in New Jersey, USA with a high abundance and diversity of flowers. These meadows were managed for pollinator-attractive, summer-blooming forbs through seed addition, and a combination of mowing, burning, and weed removal. Most flower species present in the meadows are native to the eastern United States. We collected our data during peak bloom and maximum day length (6 June to 20 August, 2016), and during good weather (sunny enough for observers to see their own shadow, no precipitation). We visited each site for three consecutive good weather days over five evenly spaced sampling rounds in the 11-week period of our study. In all analyses, we assume that bees and flowers detected at a site within one 3-day sampling round co-occurred. In contrast, we assume that turnover of both plant species in bloom and bee species activity can occur in the ~10 days between sampling rounds.

During each 3-day sampling round, an observer walked parallel transects through the meadow (which ranged in size from 0.8–2.2 ha; mean=1.4 ha), observing every open flower within a moving 1-m semicircle, and net-collecting any bee seen actively foraging, which we defined as contacting anthers or collecting nectar from a flower (Fig. S1). We collected all bee species except *Apis mellifera* L., the domesticated western honey bee, because *Apis* males do not forage. Observations began as soon as pollinator activity picked up in the morning (7–9 am) and continued into the late afternoon or evening until pollinator activity slowed substantially. Observers sampled nearly continuously, in 30-minute timed collection bouts with short breaks in between. If inclement weather precluded a minimum of six 30-minute sampling bouts in a day, we added an additional day to the sampling round as soon as weather permitted.

Flower species were identified in the field by the data collector. Bee species were identified using a dissecting microscope and published keys; Jason Gibbs (University of Manitoba), Joel Gardner (University of Manitoba), and Sam Droege (USGS) assisted with identification for bees in the genera *Andrena, Anthophora, Coelioxys, Halictus, Heriades, Hoplitis, Hylaeus, Lasioglossum, Megachile, Melissodes, Nomada, Osmia, Pseudoanthidium, Ptilothrix, Sphecodes, Stelis*, and *Triepeolus*, and at least one of them confirmed voucher specimens for every species. We determined every specimen to species except for the following four complexes: Most bees in the genus *Nomada* with bidentate mandibles (*ruficornis* group) were treated as one species. All specimens from the *Hylaeus* species complex that includes *Hylaeus affinis, H. modestus*, and at least one additional species, informally dubbed “species A,” were treated as a single species, denoted *Hylaeus affinis-modestus*, because females cannot be reliably distinguished. There is a cryptic species in the genus *Halictus* unlikely to occur in our area, *Halictus poeyi*, which is not morphologically distinct from *H. ligatus*; we treat all specimens in this complex as *Halictus ligatus*. We could not confidently separate all specimens of the two closely related *Lasioglossum* species *Lasioglossum hitchensi* and *L. weemsi*. Thus, we treat all specimens from either species as one, denoted *Lasioglossum hitchensi-weemsi*. All bee specimens are curated in the Winfree lab collection at Rutgers University, and the data used in this paper are available from the Dryad Digital Repository http://dx.doi.org/XXXXXXX (Roswell et al.)

### Analytical methods

We performed all statistical analyses and simulations using R 3.5.1 (R Core Team, 2018).

#### 1) How much do male and female bee diets overlap?

To compare the diets of male and female bees, we used the Morisita-Horn index of resource overlap (Horn, 1966; Morisita, 1959). This dissimilarity index compares the proportion of all female bees found on each flower species to the proportion of all male bees found on each flower species. In other words, it compares the contribution of each flower species to female diets (where this term includes the food that females collect for themselves and also to feed to young) to the contribution of the same flower species to male diets. The Morisita-Horn index ranges from zero (completely similar) to one (maximally dissimilar), and has several good properties for our purposes. First, it uses proportions, placing visits from male and female bees on the same scale, even though most visits come from females. Second, it is much more sensitive to large proportions than to small ones, thereby down-weighting the contribution of flower species for which we have little information. Third, the Morisita-Horn estimates are resilient to undersampling and uneven sample size between groups (Barwell, Isaac, & Kunin, 2015).

To determine whether the male-female differences we observed exceeded those expected by chance, we compared the observed compositional dissimilarity between flower visits from male and from female bees to dissimilarity measures from a null model that randomly permuted the bee sex associated with each flower-visit record. This permutation holds constant the total number of male and of female visits, and the total number of visits to each flower species from both sexes combined (Fig. S2). The range of dissimilarity values from this simulation is the difference we would observe in our sample, if there were no true difference in flower species use between males and females of the same bee species. We evaluated the hypothesis that male and female diets overlap less than would be expected by chance; thus we use a one-sided alpha of p<0.05. We iterated this null simulation 9999 times, which was sufficient to stabilize p-values near our chosen alpha (North, Curtis & Sham 2002). When the observed dissimilarity was greater than 9500 of the 9999 simulated dissimilarities, we concluded that we had detected a difference in the pattern of floral visitation between conspecific male and female bees, given the observed diet breadth and abundance of each sex.

To compare the diet overlap we observed between sexes to a meaningful benchmark, interspecific diet overlap, we repeated the same null model analysis, this time comparing females of the focal species to females of other species. We performed one analysis for each bee species for which we collected at least 20 visitation records for each sex (19 species). This sample size threshold is arbitrary, but null model variance shrinks with sample size, such that apparent patterns for species with smaller sample sizes are rarely interpretable (Fig. S3). Because we analyze 19 bee species, females of each species are compared to 18 others. We then compared the male-female difference (observed minus mean null dissimilarity in flower communities visited) to the analogous species-species difference (observed minus null dissimilarity).

For this analysis, which evaluates holistic differences between male and female bees of the same species, we combined observations across the full season and all sites. This allows us to observe foraging niche differences that are driven by flower and/ or bee phenology, in addition to any sex-specific floral preference.

#### 2) To what degree are particular flower species disproportionately visited by bees of one sex?

This analysis uses our entire data set of 153 bee species to determine whether particular flower species are disproportionately visited by male or female bees, and whether the answer varies by bee species. We can infer a preference difference between male and female bees for a flower species when predicted odds of visitors to that flower species being male are especially high or low. To do this, we use a random effects model in which bee sex is the response, and flower species, bee species, site, and their interactions are random effects. We statistically control for variation in the overall sex ratio across bee species through a random intercept of bee species, and variation in sex ratios across sites, through random intercepts for site, and the site-bee species and site-flower species interactions. Because it is unlikely that, within bee species, sex ratios at birth vary greatly across space, any variability attributed to site terms would likely result from differential overlap of bee foraging activity and flower bloom across space.

We call this model the “summed model” because we sum interactions observed across the entire season (all five sampling rounds) at each site. In the summed model, the relationship between phenological overlap and the odds of flower-visiting bees being male would be incorporated into the species effects. This perspective is helpful for considering flower species’ contributions to the overall diets of male versus female bees. We fit the model with the R package lme4 (Bates, Maechler, & Walker, 2016) with the following call:

~~~
***Summed model***
lme4∷glmer(bee_sex ~ (1|site)+ (1|flower_species)+ (1|bee_species)+ 
        (1|flower_species:bee_species)+ (1|site:bee_species)+ 
        (1|site:flower_species), family=”binomial”, data=data)
~~~

We included bee species and site as random, rather than fixed, effects to directly compare the variability in bee sex associated with each of these predictors to the variability associated with flower species (preference). Comparing the overall variability across these groups was more important to us than assessing predictions on a per-site or per-bee-species basis. We fit flower species, the primary covariate of interest, as a random effect to facilitate model fitting (fewer degrees of freedom) as well as interpretation. In our summed model, we included all two-way interactions, but omitted the three-way interaction, bee species by flower species by site. Although the sort of context-dependent preference this term could represent (e.g. males from bee species *1* prefer flower *A* at one site (relative to females), but shun it at another) may exist in nature, it is unlikely we would estimate it accurately in our model.

We confirmed model convergence by comparing several fitting methods using the allFit function in lme4 (Bates et al., 2016), which all showed similar parameter estimates (Table S1). We tested whether residuals from our model fit were overdispersed using Bolker’s function “overdisp” (Bolker, 2017), and visually assessed our additivity assumptions with binned residual plots (Gelman & Hill, 2007) (Fig. S4).

#### 3) To what extent are differences in floral use driven by preference, rather than phenological differences between male and female bees?

Over the 11 weeks of our study, we observed turnover in bee species, in flower bloom, and within-bee species changes in sex ratio. Therefore, phenological overlap between male versus female bees and the bloom period of particular flower species, rather than preference of those bees for those flowers, may explain much of the variation in sex ratio we observed across visitors. In question 3, we are explicitly interested in distinguishing sex-specific diet *preferences* from variable *use* resulting from seasonal resource availability and male vs. female abundance. We do this in the “seasonal model” by incorporating sampling round (our measure of phenology) as an additional random intercept effect, along with random intercepts for the interactions between sampling round and the other covariates. We chose to include sampling round as a random effect because this enables direct comparison to all other terms in both models. We ignored the three-and four-way interactions between bee species, flower species, and other covariates. We fit this model with the following call in the R package lme4, with new terms in bold:

~~~
**Seasonal model**
glmer(bee_sex ~ (1|site)+ (1|flower_species)+ (1|bee_species)+
        (1|flower_species:bee_species)+ (1|site:bee_species)**+ 
        (1|site:flower_species)+ (1|sampling_round)+ (1|site:sampling_round)+ 
        (1|flower_species:sampling_round)+ 
        (1| bee_species:sampling_round)+ 
        (1|site:bee_species:sampling_round)+ 
        (1|site:flower_species:sampling_round)**, family=”binomial”, data=data)
~~~

Our index of preference for both the *summed model* and the *seasonal model* is the change in odds that a bee is male when the flower species it visits is given. To describe the importance of model terms, we calculated a bootstrapped median odds ratio using code from Seth (Seth, 2017), which gives the expected difference in odds that a flower-visiting bee is male between levels of a predictor (Merlo et al., 2006). For example, a median odds ratio of five for the main effect of sampling round would indicate that the odds of a flower-visiting bee being male differ by about a factor of five between sampling rounds, while a median odds ratio of one would indicate that the odds of a flower-visiting bee being male do not change across rounds. If the median odds ratio is large for flower species in both models, we could say that there are intrinsic (i.e. not simply phenological) properties of flower species identity that male or female bees prefer. If flower species is a strong predictor of bee sex in the summed model but not in the seasonal one, we would still conclude that flower species often contribute more strongly to the diet of one sex than the other, though these differences may not arise due to differing preferences. If the sampling round terms have large median odds ratios, then accounting for phenology is critical for identifying differences in preference in addition to differences in use.

## Results

In total we collected 18,698 bee specimens belonging to 152 bee species (Table S2) from a total of 109 flower species (Table S3), which together comprised 1417 unique species-species interactions. Roughly 18% of specimens were male (n=3372). Thus, the overall ratio of male to female bees we collected was 0.22, although this ratio varied markedly between flower species (Fig. 1).

**Figure 1.**
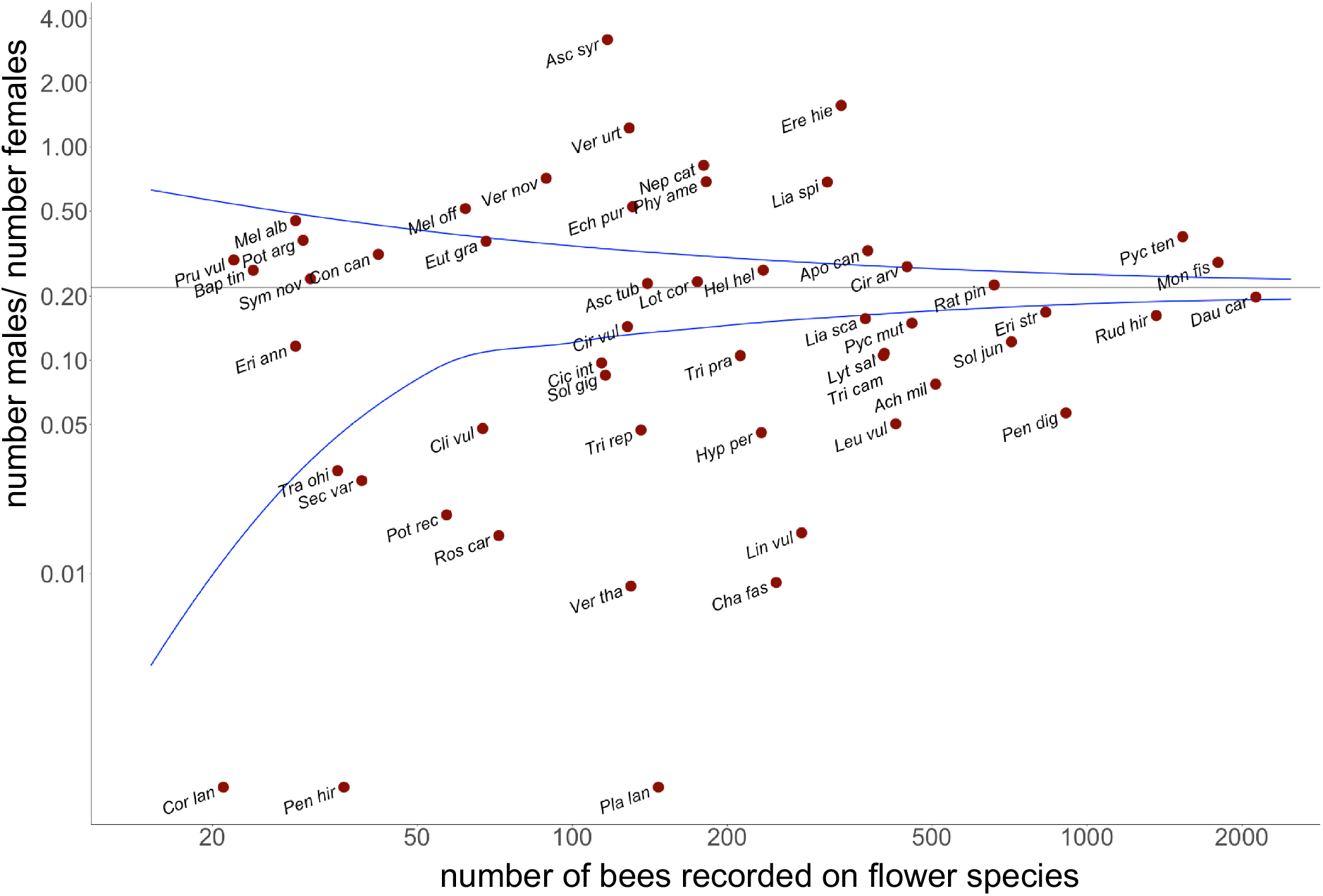
The sex ratio (M:F) of flower-visiting bees varies across flower species. Each red point represents a flower species, the first three letters of the Latin genus and species names for the flower species label each point. The x-axis is the number of bees collected from that species, the y-axis is the ratio of male to female bees collected from the flower. Flower species that received >19 visits are plotted (n=54). Blue lines are smooth fits to the 97.5^th^ and 2.5^th^ percentiles of the binomial distribution given by the observed ratio of males to females in our overall dataset (M/F=0.22; i.e. M/(M+F)=0.18). This distribution represents our expectation for random variation in sex ratio across flower species, if the sex ratio of flower-visiting bees is independent of flower species identity (male and female bees exhibit the same floral preferences), and remains nearly constant across time and space.

### How much do male and female bee diets overlap?

We found that male and female bee diets overlap significantly less than would be expected at random (Fig. 2), and that the differences in diet composition between male and female bees of several species were of similar magnitude to the differences in diet between species of bee (Fig. 3).

**Figure 2:**
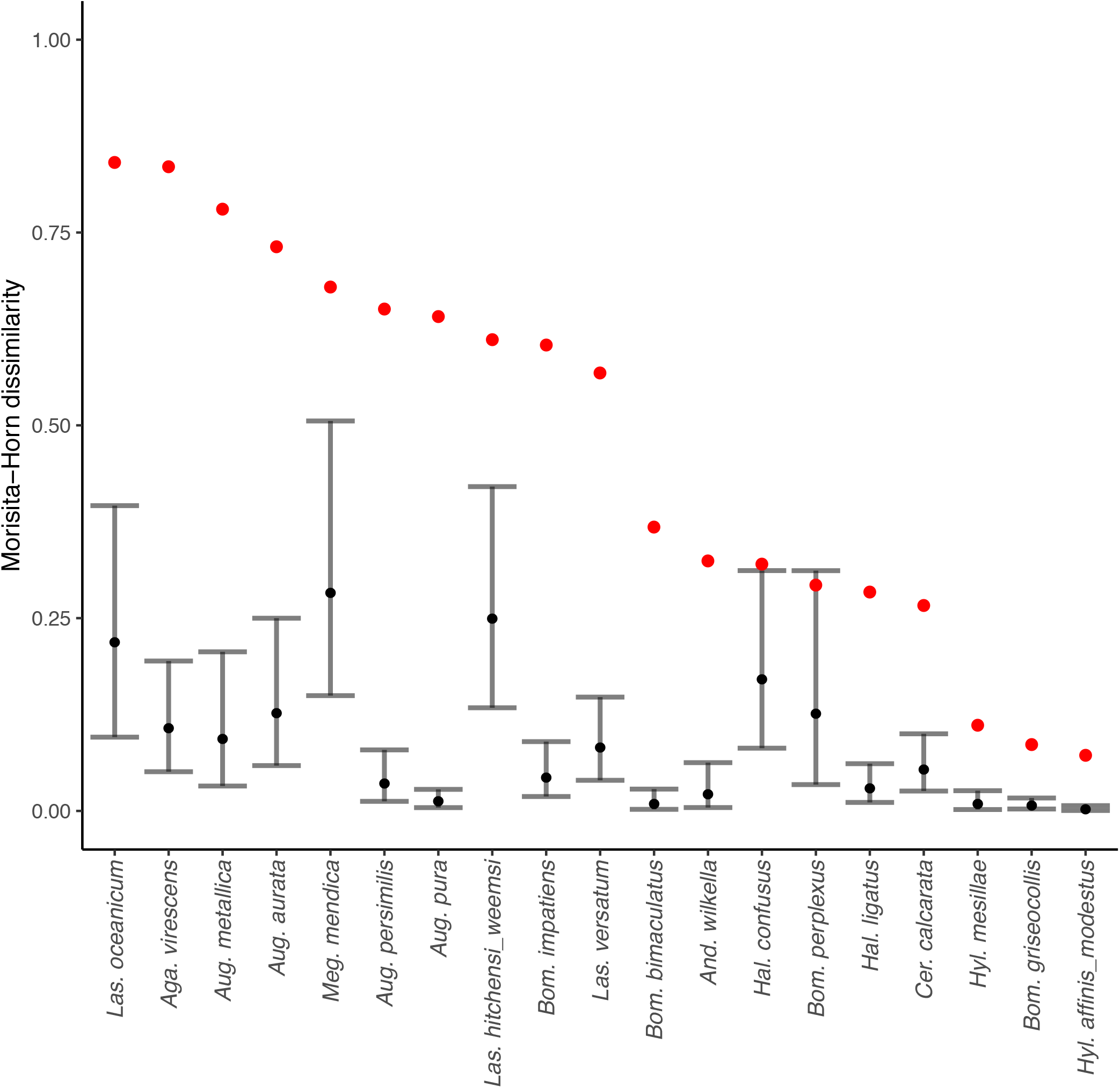
Flower visit patterns of male and female bees of the same species differed significantly. Red points are observed Morisita-Horn dissimilarities between flower communities visited by all male and all female bees of a particular species across all sites and sampling rounds. Black points are the mean dissimilarity (gray bars, 95% CI) from a permutation-based null model that randomly shuffles the sex associated with each visit record, maintaining the total number of males, females, and overall combined visits to each floral species.

**Figure 3.**
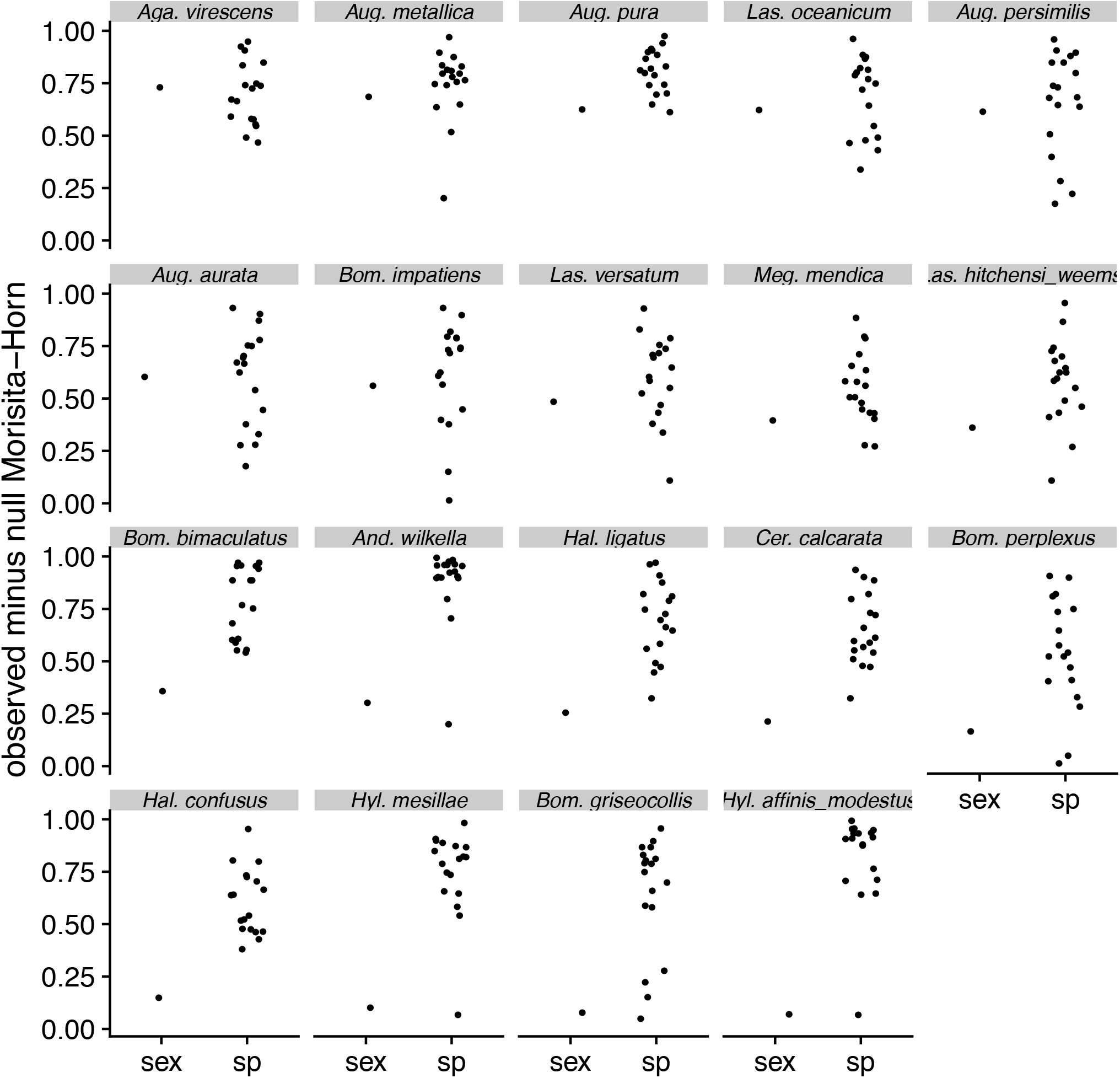
The diets of male and female bees of the same species can be as dissimilar as the diets of females of two different bee species. Dissimilarities in this figure are the observed statistic minus, for each pairwise comparison, the mean dissimilarity in the null model. Each panel focuses on a bee species (panel name) and shows: above the label “sex”, observed diet dissimilarity between male and female bees of the focal species, minus the average null dissimilarity resulting from randomly permuting the sex identity of each visit record; above the label “sp”, observed diet dissimilarity between female bees of the focal species and each other bee species, minus the average null dissimilarity resulting from randomly permuting the species identity of each visit record.

### To what degree are particular flower species disproportionately visited by bees of one sex?

The sex ratio of flower-visiting bees varied across species of flower (Fig. 2). After controlling for bee species identity (the strongest predictor of sex in our models, Fig. 4), and site, we still found that some flower species received a disproportionate number of male bee visitors (Figs 4–5). The median odds ratio for the main effect of flower species was 3.6 (bootstrapped CI 3.0–4.2) in our summed model, indicating that, typically, the visitor sex ratio differs between two flower species by more than a factor of 3. Furthermore, we observed sex-based differences in flower use specific to particular bee species: the median odds ratio for the flower species by bee species interaction in our summed model was nearly as large (median=3.1, bootstrapped CI 3.0–3.3) as the main effect of flower species. By contrast, sex ratios are not expected to differ between sites (median odds ratio for main effect of site=1).

**Figure 4.**
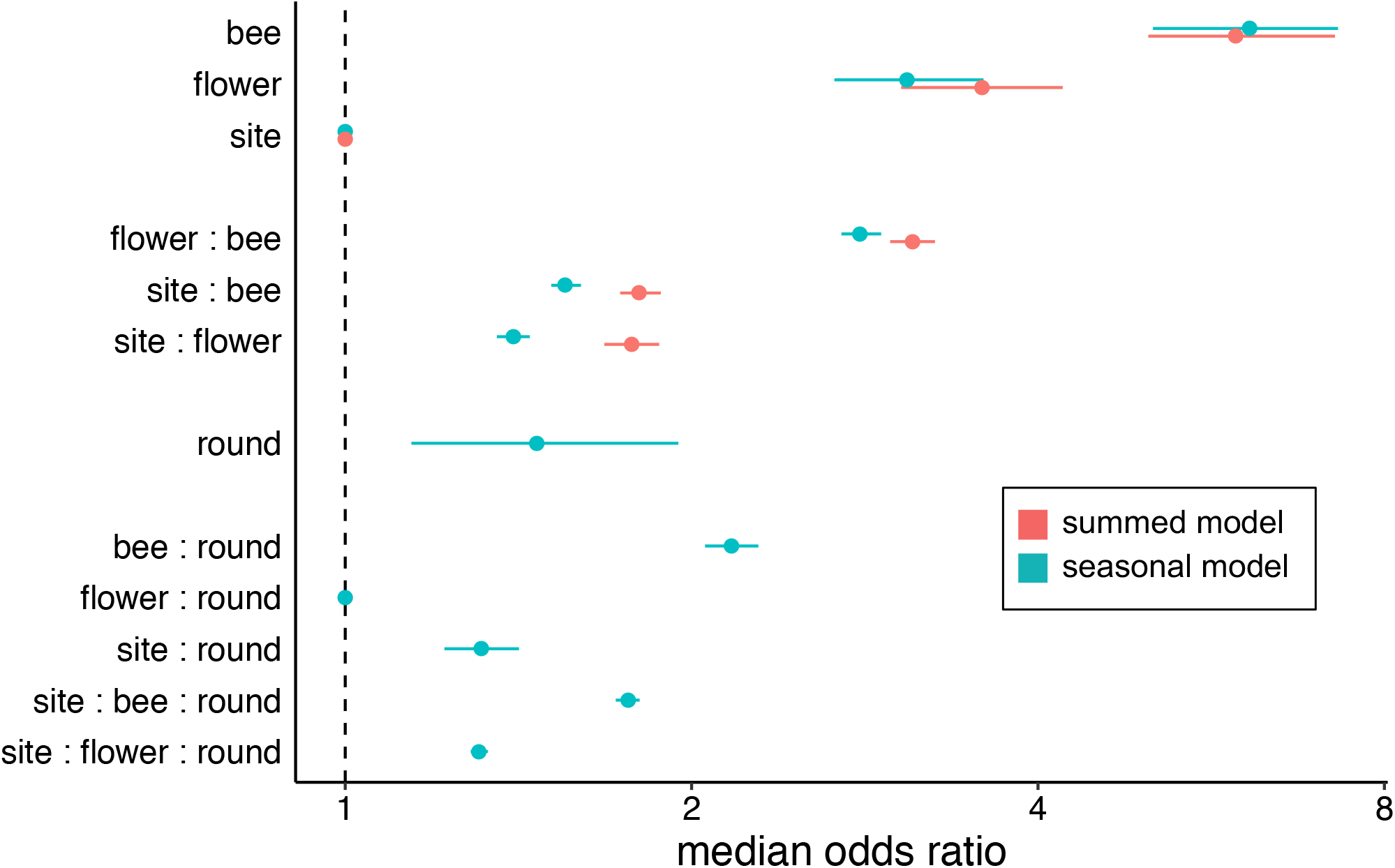
Flower species, along with bee species, predicts the sex of visiting bees, which suggests floral preferences differ between male and female bees. Flower species is an important predictor of bee sex even after accounting for phenology (seasonal model). For each term (“bee”= bee species, “flower”=flower species, “round”=sampling round) in each model, the median odds ratio (+/- 95% bootstrapped credible interval) indicates the expected difference in odds that a flower-visiting bee is male between two levels. For example, a median odds ratio of 3.7 for the flower species term means the odds of a visitor being male are expected to differ by a factor of 3.7 between two randomly selected species of flower.

**Figure 5.**
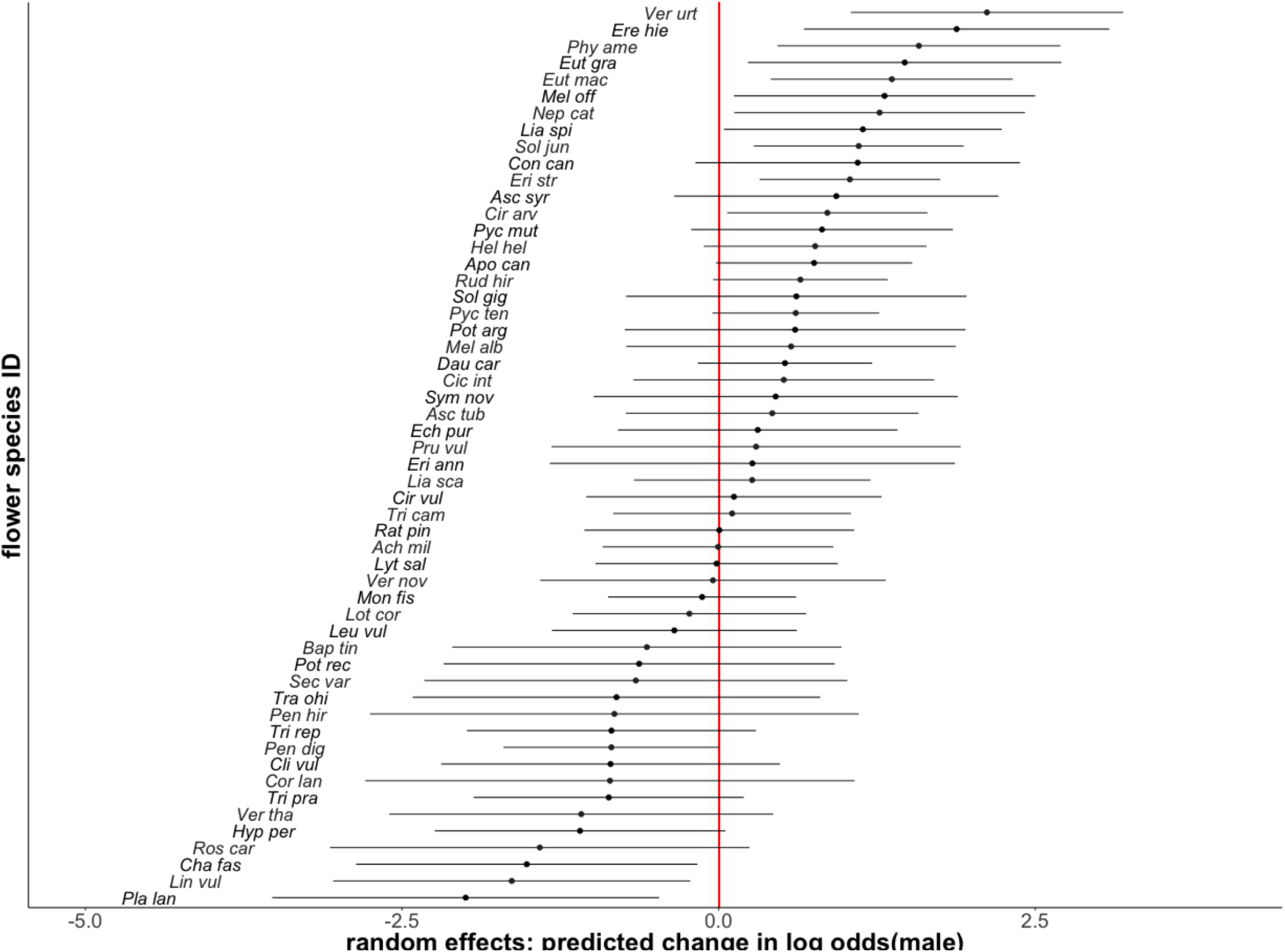
Male bee preferences for and against flower species vary across flower species. Each point is the conditional mode of the random effects prediction (the random-effects analog to an estimate), for a flower species that received at least 20 visits, on the logit scale. Zero represents the odds of a visitor being male on a random flower, and -2 or 2 indicates a ~7 fold decrease or increase in those odds, given flower species identity. Error bars are the square root of the conditional variances on the conditional mode * 1.96, and can be interpreted as the expected range in which the random effect for a particular flower truly lies, analogous to 1.96 times the standard error of the mean for a fixed effect.

### To what extent are differences in floral use driven by preference, rather than phenological differences between male and female bees?

The flower species blooming in our system turned over throughout our 11-week sampling period, with several highly visited species blooming for only one of the three months during which we sampled. This turnover, along with potential sex-specific bee flight seasons, means that differences in diet between male and female bees could reflect seasonal availability and use, without also indicating preference differences between the sexes. Indeed, phenology predicts bee sex somewhat, with the odds of a flower-visiting bee being male expected to change by a factor of 1.5 (bootstrapped CI 1.1–1.9) between sampling rounds (Fig. 4). Phenological patterns of male vs. female flight seasons vary across bee species; the median odds ratio for the bee species by sampling round interaction is 2.2 (bootstrapped CI 2.1–2.3) (Fig. 4). Even after accounting for these effects, however, there remains a strong association between the species of flower a bee visits and its sex (Figs 4–5). The relative effects of each flower species on the sex of its visitors were changed very little by accounting for phenology; Pearson and Spearman correlations between the random effect of flower species in the seasonal model and the same random effect in the simpler summed model were both 0.98. In addition to finding overall preference difference between male and females, we found evidence for bee-species-specific difference in floral preferences between the sexes (median odds ratios in both models for the bee species by flower species interaction > 2.8).

## Discussion

We found strong differences between the flower species preferences of male and female bees. The difference in floral visits between male and female bees of the same species was similar in magnitude to differences between females of different species. The partitioning of the floral community among bee species is a primary focus of pollination ecology and ecological network analysis (Bascompte & Jordano, 2014), but male bees are typically disregarded or lumped together with their female counterparts. Our study suggests this may represent an important oversight. Further, our study provides strong confirmation of the few studies that investigate the foraging behavior of male bees, which found that males play a unique role in plant pollination (Cane, 2002; Cane et al., 2011; Ogilvie & Thomson, 2015; Pascarella, 2010). Our result also implies that male bees contribute substantially to the complexity of plant-pollinator networks in nature, and that network analyses might benefit from separating males and females into different nodes (Bolnick et al., 2011; Zwolak, 2018).

Phenology, a previously reported mechanism for distinct use of floral resources by male and female bees (Ogilvie & Thomson, 2015; Robertson, 1925), explained some variation in the sex ratio of flower-visiting bees, but was less important than flower species identity over the period of our study. We expected to find an effect of phenology because both the identity of the flower species blooming within sites, and also the sex ratio of foragers within bee species, vary across the season. Males emerge first in most solitary bees; for social species, initial broods usually consist primarily of female workers, then males and reproductive females emerge at the end of the colony cycle (P. G. Willmer & Stone, 2004). Surprisingly, however, phenology only weakly predicted the sex of flower-visiting bees. This is despite the fact that, as predicted by natural history, the sampling round(s) in which males were relatively more prevalent depended on bee species (the bee species by sampling round interaction was much bigger than the sampling round main effect; Fig. 4). This indicates that our evidence for floral preference differences between male and female bees was robust to accounting for seasonal turnover in flower species bloom, bee species flight seasons, and the sex ratios within bee species.

Whereas female bees collect both nectar and pollen, male bees forage primarily for nectar to fuel flight. Thus, we predicted that male bees would avoid flowers that produce no nectar. Indeed, in both our models, the predicted odds of a bee visiting a nectar-less flower species being male were approximately half that of a bee visiting a flower species that produces nectar (Fig. S5). A second biological difference between male and female bees is that adult male bee activities orient around mate seeking (Alcock et al., 1978). These behaviors, such as patrolling routes (Barrows, 1976) or seeking flowers visited by conspecific females (Rossi, Nonacs, & Pitts-Singer, 2010) could generate differences from females via complementarity (males visiting flower species not visited by females), or nestedness (one sex primarily visiting a subset of species visited by the other). We found evidence for both (Fig. S6). Divergent floral preferences between sexes may reflect nutritional needs or mating behavior, or simply biases resulting from previous flower encounters, or visual or olfactory sensitivities that differ between the sexes (Robert, Frasnelli, Collett, & de Ibarra, 2016; Somanathan, Borges, Warrant, & Kelber, 2017; Streinzer, Kelber, Pfabigan, Kleineidam, & Spaethe, 2013).

While natural and semi-natural habitats are critical landscape elements for many bee species (Senapathi, Goddard, Kunin, & Baldock, 2016), what constitute suitable and/ or limiting resources within these habitats remains less clear (De Palma et al., 2015). Flowers, which provide food for adult and larval bees, are likely among them (Roulston & Goodell, 2011). Whether floral diversity *per se* tends to benefit individual pollinator taxa remains unclear (Spiesman, Bennett, Isaacs, & Gratton, 2017; Sutter, Jeanneret, Bartual, Bocci, & Albrecht, 2017). However, complementary flower species use between the sexes implies a mechanism by which a bee species could benefit from a diversity of flower choices. In addition, it is currently unknown how the distinct foraging niches of male bees mediate either the robustness of pollinator communities to species loss and environmental perturbations (Brosi & Briggs, 2013; Ramos-Jiliberto, Valdovinos, Moisset de Espanés, & Flores, 2012; Tur et al., 2014), or the effectiveness of different habitat ameliorations (Rundlöf, Persson, Smith, & Bommarco, 2014; Rusterholtz & Erhardt, 2000; Williams & Lonsdorf, 2018). This study suggests that both questions warrant further investigation.

Patterns in bee-flower interaction data can arise from the sampling process itself (Blüthgen, 2010; Fründ, McCann, & Williams, 2016). Our analyses control for these patterns. To evaluate diet overlap, we used a dissimilarity index that downweights rare diet items, and implemented a null model that accounts for differences that could arise from sampling and abundance effects. To evaluate preference, we used random effects models that incorporated all (nearly 19,000) observations, and shrank extreme values for rarely observed species-species interactions towards the global mean for each effect. Thus, our estimates for sex-specific preferences should be robust to the inevitable under-sampling of rarer taxa. Establishing differences in preference between categories of bees such as males and females, even when resource availability is seasonal and difficult to quantify, is possible using methods such as these, though absolute preference remains elusive.

Pollination ecology and pollinator conservation still face the question of how important sexually dimorphic foraging is. Does it enhance or reduce the stability of bee populations? Should pollinator restorations explicitly include “male bee” flowers and “female bee” flowers? Are floral traits under selection to favor female versus male visitors? While our study does not answer these questions, by showing that the diets and preferences of male bees commonly differ from those of their female conspecifics, we suggest they are worthy of future study.

## Author contributions

MR and RW conceived the ideas and designed field methodology; MR, JD, and RW designed statistical methodology. MR collected the data; MR, RW, and JD analyzed the data; MR and RW led manuscript drafting. All authors contributed critically to the drafts and gave final approval for publication.

## Acknowledgements

We thank Daniel Cariveau and Neal Williams for study design suggestions, and Jason Gibbs, Tina Harrison, and Dylan Simpson for comments on an earlier draft. A USDA Conservation Innovation Grant to the Xerces Society for Invertebrate Biology (lead PI) and RW (senior investigator) funded data collection. MR was supported by an NSF Graduate Research Fellowship. Cameron Kanterman and Riva Letchinger helped collect field data; Kurtis Himmler, Tiffany Bennet, and professional taxonomists Jason Gibbs, Joel Gardiner, and Sam Droege helped with bee species determinations. Mercer County, the Institute for Advanced Study, Somerset County, and the Raritan Headwaters Association provided site permission.

---

**Figure S1:**
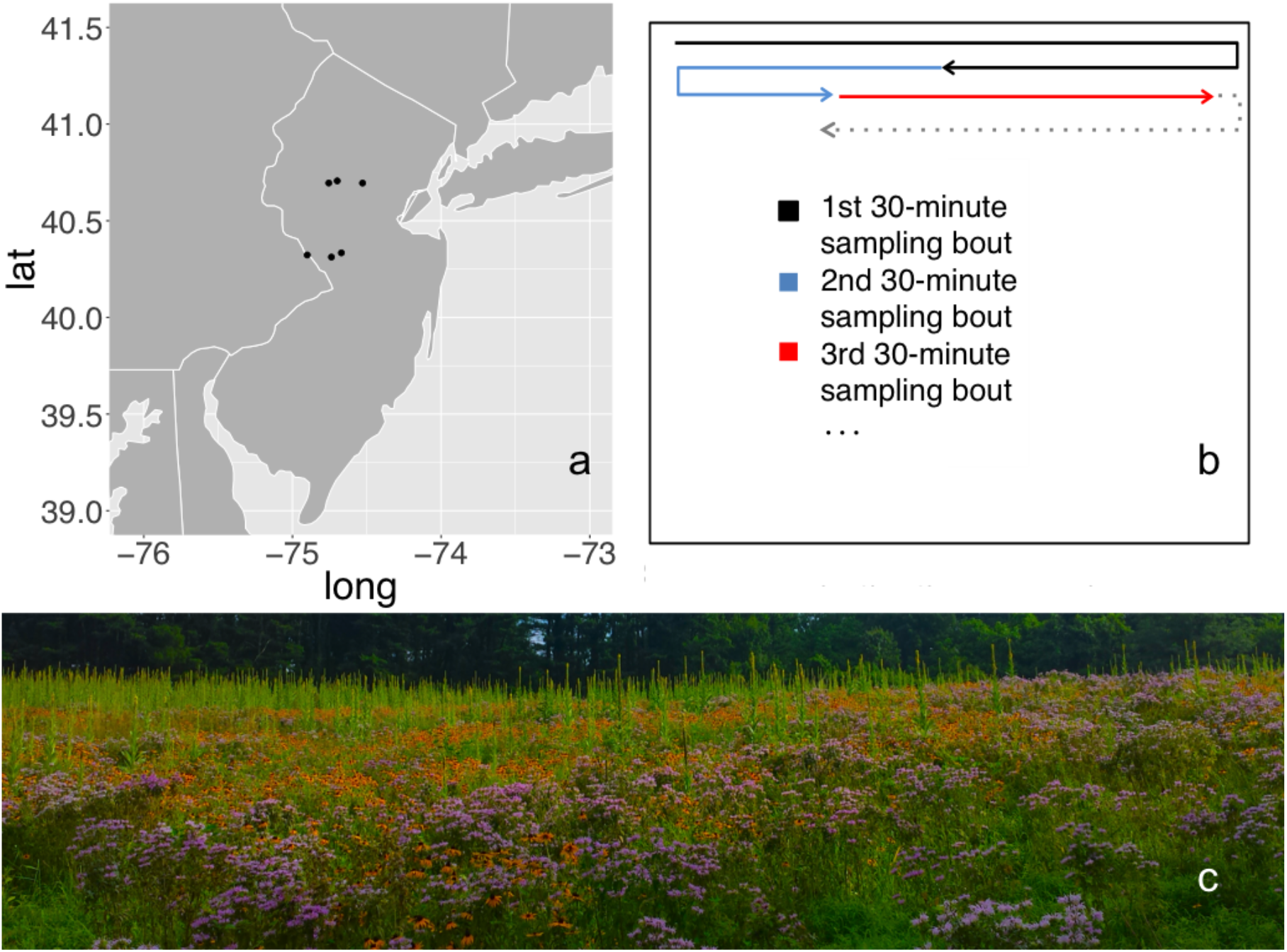
Sampling scheme. (a) The six study sites in central New Jersey, USA. (b) Schematic sampling diagram (not to scale). One observer walked parallel 2m transects covering the entire sampling area. Each 30-minute sampling bout resumed where the previous one left off; observers typically covered the entire meadow once over a 3-day sampling round. (c) The southwestern-most site in peak bloom.

**Figure S2.**
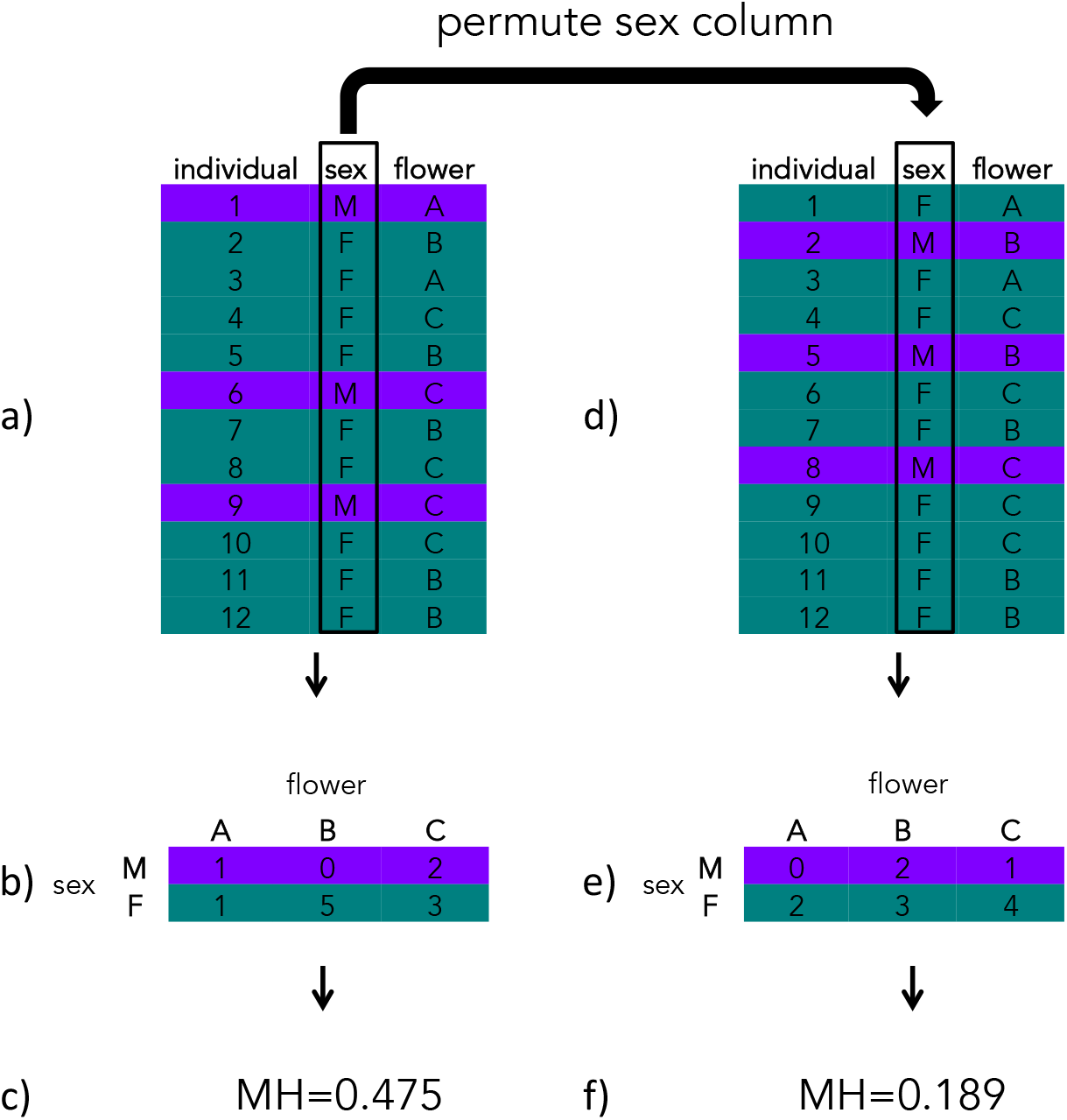
Schematic cartoon of our simulation for the dissimilarity values associated with our null hypothesis that diets of male and female bees do not differ. (a) Each collection record for each bee species associates the sex of an individual bee to the flower species from which it was collected. (b) To compute the dissimilarity between males and females, we compare all visits to each flower species from males (purple vector) to all visits to each flower species from females (green vector). (c) The Morisita-Horn index summarizes the differences between the two vectors as a value between 0 (identical) and 1 (maximally dissimilar). (d) For our null model, we shuffle the sex column from our observation table. (e) This produces two null vectors. The row and column sums for the matrices in (b) and (c) are identical, but the elements can differ. (f) For our null model, we compute the dissimilarity between the null vectors. We repeated steps d-f 9999 times to generate confidence intervals for the null hypothesis that the sex of a visiting bee is unrelated to the flower species it is collected from. When comparing the flower species visited by different species of bee, we conducted an analysis identical except that rather than comparing two sexes of the same species, we compared two species of the same sex (i.e. exchanging “sex” and “species” throughout figure S1).

**Figure S3.**
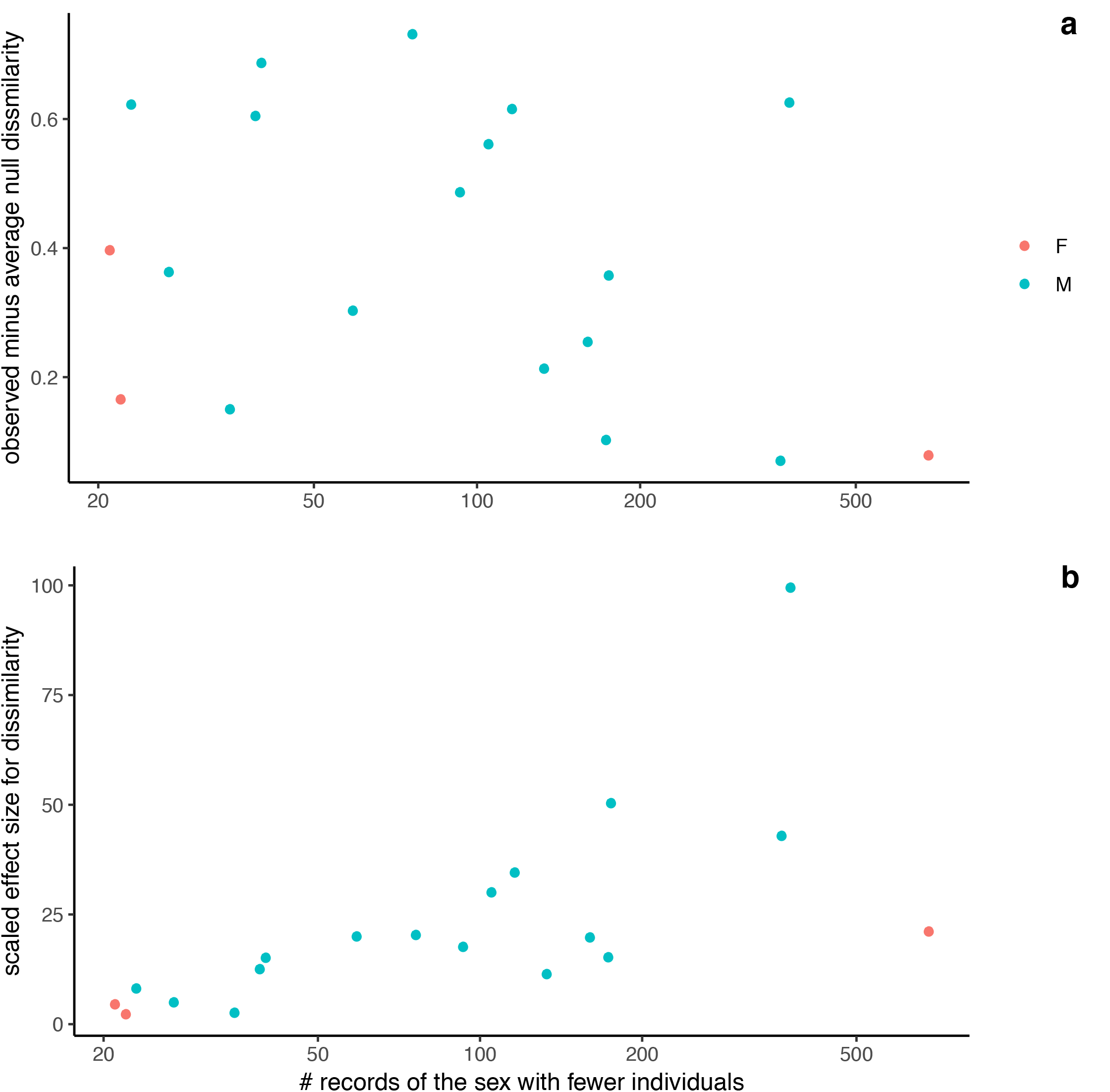
Effect size for diet dissimilarity is independent of sample size, while standardized effect is strongly driven by the number of individuals of the sex with the fewest records. a) Observed Morisita-Horn dissimilarity in flower communities visited by male and female bees of a single species, minus average null dissimilarity vs. the number of records for the less frequently observed sex. b) Observed minus null dissimilarity in composition of flowers visited by male and female bees of a single species, scaled by the variation in the null model, versus the number of records for the less frequently observed sex.

**Figure S4.**
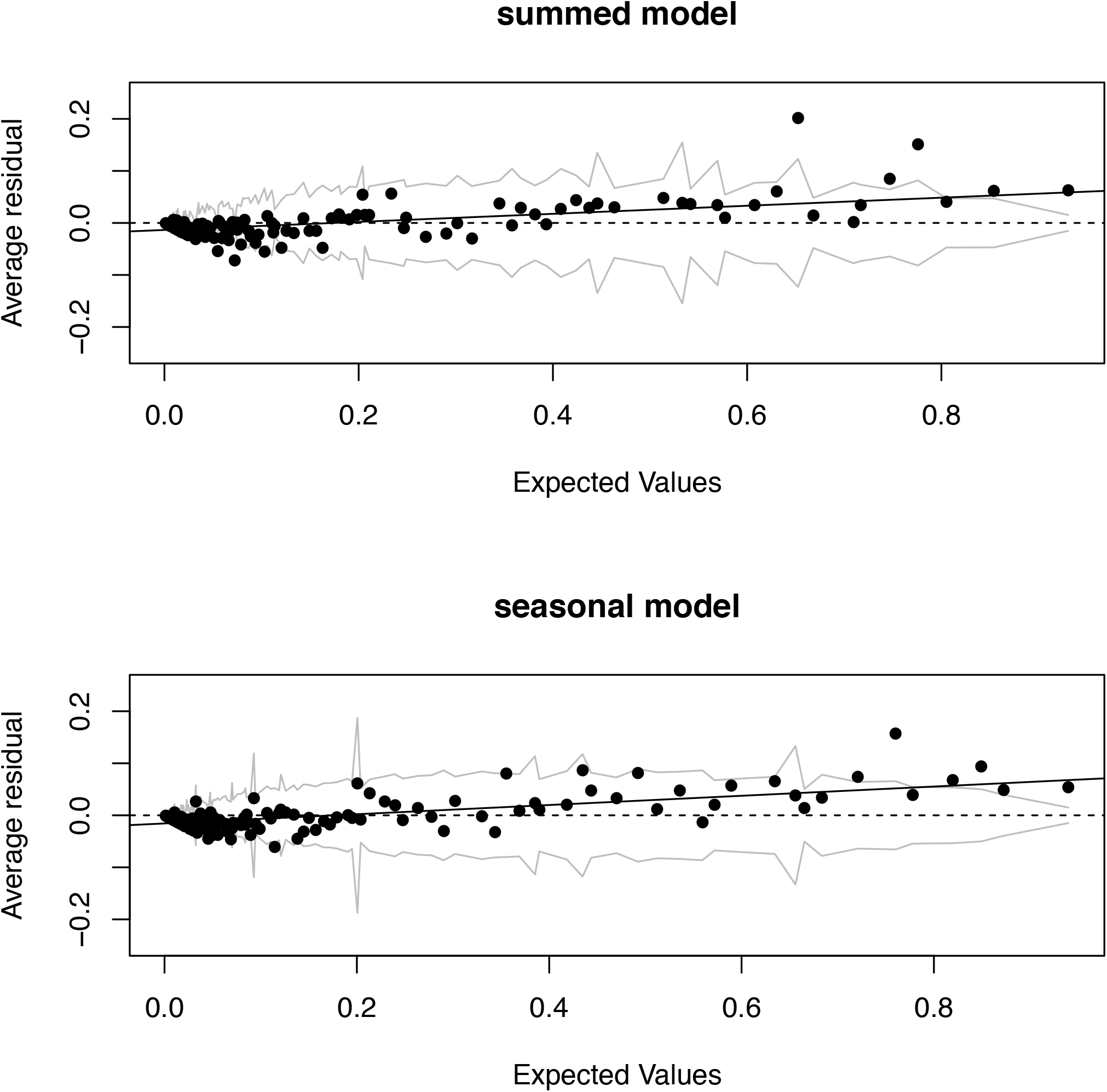
Binned residual plots for each model show minor violation of the additivity assumption. Residuals and predicted values on the probability scale.

**Figure S5.**
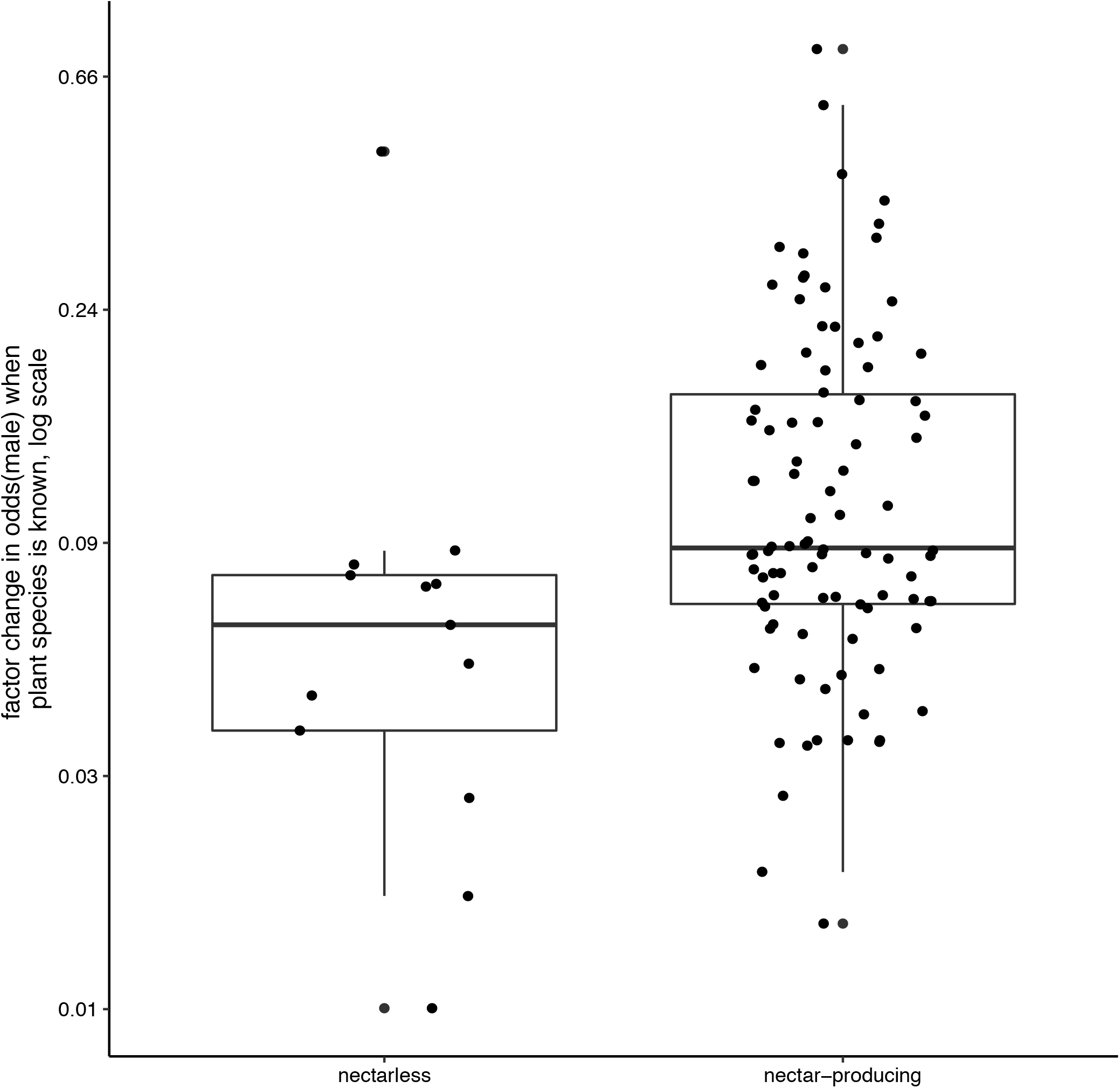
Seasonal model predictions are consistent with the hypothesis that male bees avoid flower species that do not produce nectar, relative to females. Each point is the random effect prediction (change in odds that a bee visiting that flower is male) for a flower species.

**Figure S6.**
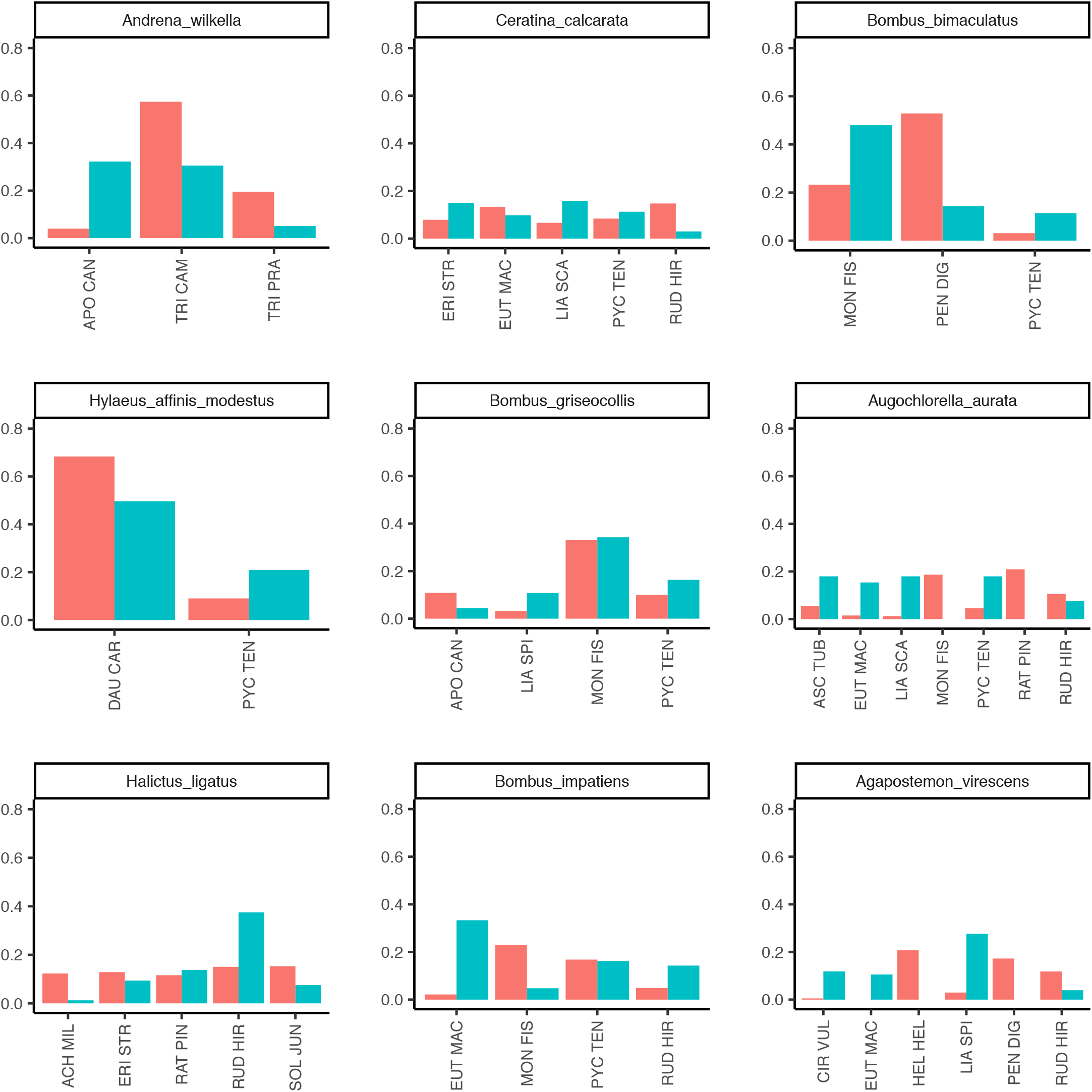
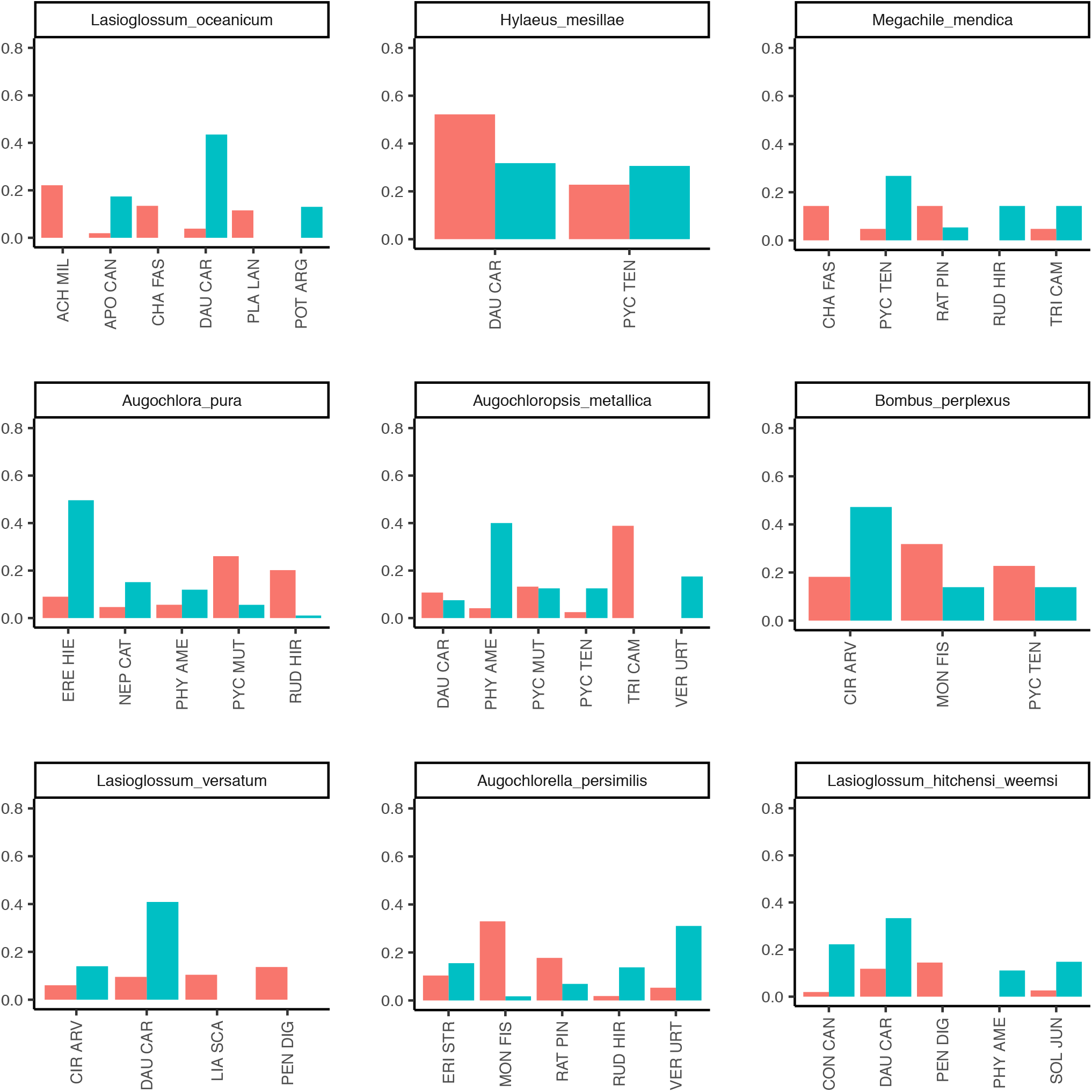
Dissimilarity in flower communities visited by male and female bees arise due to complementarity in addition to nestedness patterns. For each bee species, the proportion of male (blue) and female (red) visits to each flower species that received >10% of at least one sex’s visits are pictured. Due to omitted flower species, bars may sum to <1.

**Table S1.**
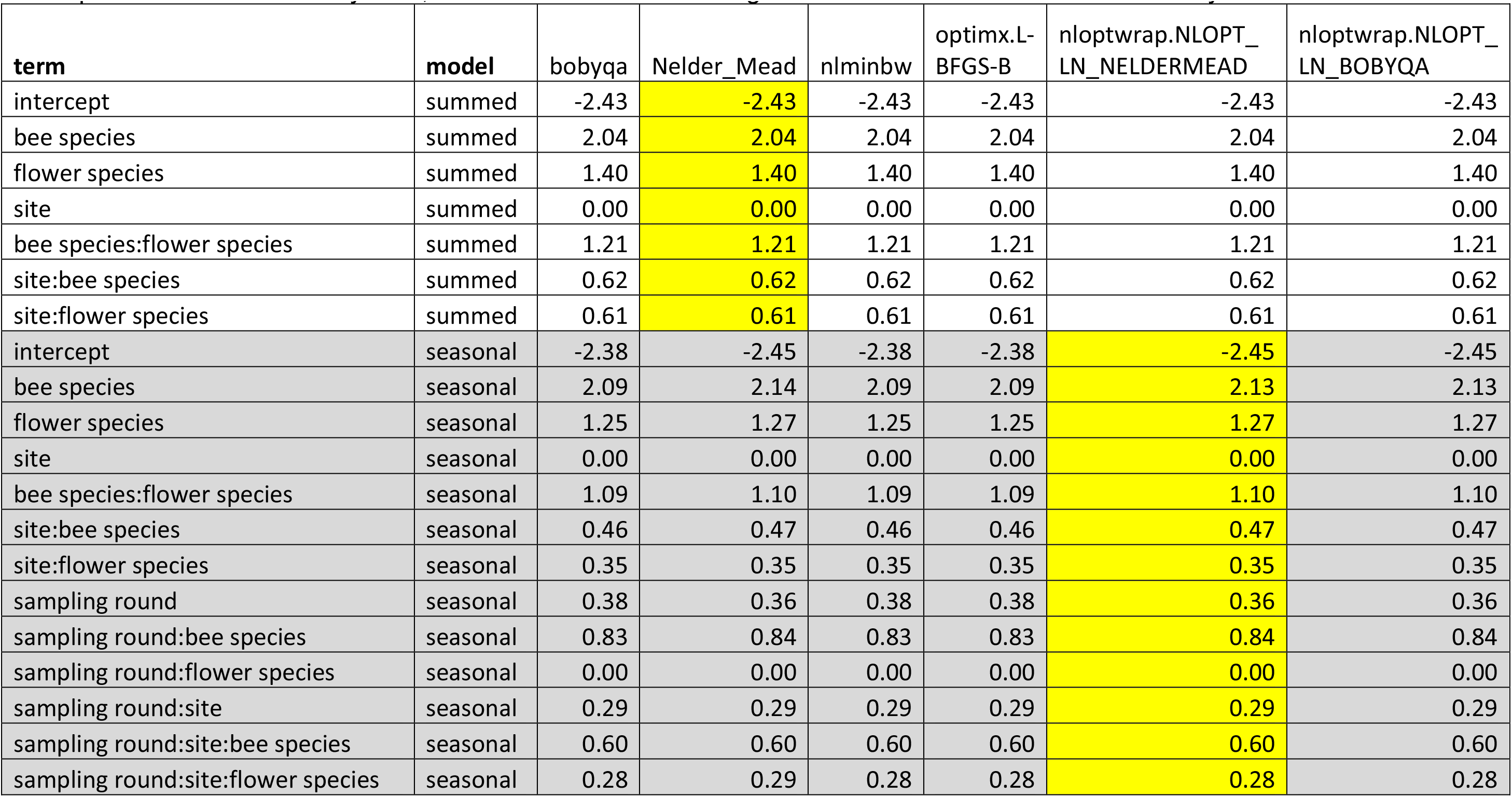
Model convergence confirmed based on similar parameter estimates across fitting routines. For each model, the estimate for each term is given for each of 6 fitting algorithms in the R package lme4. Subsequent analyses used parameter estimates in yellow, in both cases tied, for the highest estimated likelihood, with other very similar fits.

| term | model | bobyqa | Nelder_Mead | nlminbw | optimx.L-BFGS-B | nloptwrap.NLOPT_LN_NELDERMEAD | nloptwrap.NLOPT_LN_BOBYQA |
| --- | --- | --- | --- | --- | --- | --- | --- |
| intercept | summed | -2.43 | -2.43 | -2.43 | -2.43 | -2.43 | -2.43 |
| bee species | summed | 2.04 | 2.04 | 2.04 | 2.04 | 2.04 | 2.04 |
| flower species | summed | 1.40 | 1.40 | 1.40 | 1.40 | 1.40 | 1.40 |
| site | summed | 0.00 | 0.00 | 0.00 | 0.00 | 0.00 | 0.00 |
| bee species:flower species | summed | 1.21 | 1.21 | 1.21 | 1.21 | 1.21 | 1.21 |
| site:bee species | summed | 0.62 | 0.62 | 0.62 | 0.62 | 0.62 | 0.62 |
| site:flower species | summed | 0.61 | 0.61 | 0.61 | 0.61 | 0.61 | 0.61 |
| intercept | seasonal | -2.38 | -2.45 | -2.38 | -2.38 | -2.45 | -2.45 |
| bee species | seasonal | 2.09 | 2.14 | 2.09 | 2.09 | 2.13 | 2.13 |
| flower species | seasonal | 1.25 | 1.27 | 1.25 | 1.25 | 1.27 | 1.27 |
| site | seasonal | 0.00 | 0.00 | 0.00 | 0.00 | 0.00 | 0.00 |
| bee species:flower species | seasonal | 1.09 | 1.10 | 1.09 | 1.09 | 1.10 | 1.10 |
| site:bee species | seasonal | 0.46 | 0.47 | 0.46 | 0.46 | 0.47 | 0.47 |
| site:flower species | seasonal | 0.35 | 0.35 | 0.35 | 0.35 | 0.35 | 0.35 |
| sampling round | seasonal | 0.38 | 0.36 | 0.38 | 0.38 | 0.36 | 0.36 |
| sampling round:bee species | seasonal | 0.83 | 0.84 | 0.83 | 0.83 | 0.84 | 0.84 |
| sampling round:flower species | seasonal | 0.00 | 0.00 | 0.00 | 0.00 | 0.00 | 0.00 |
| sampling round:site | seasonal | 0.29 | 0.29 | 0.29 | 0.29 | 0.29 | 0.29 |
| sampling round:site:bee species | seasonal | 0.60 | 0.60 | 0.60 | 0.60 | 0.60 | 0.60 |
| sampling round:site:flower species | seasonal | 0.28 | 0.29 | 0.28 | 0.28 | 0.28 | 0.28 |

**Table S2.**
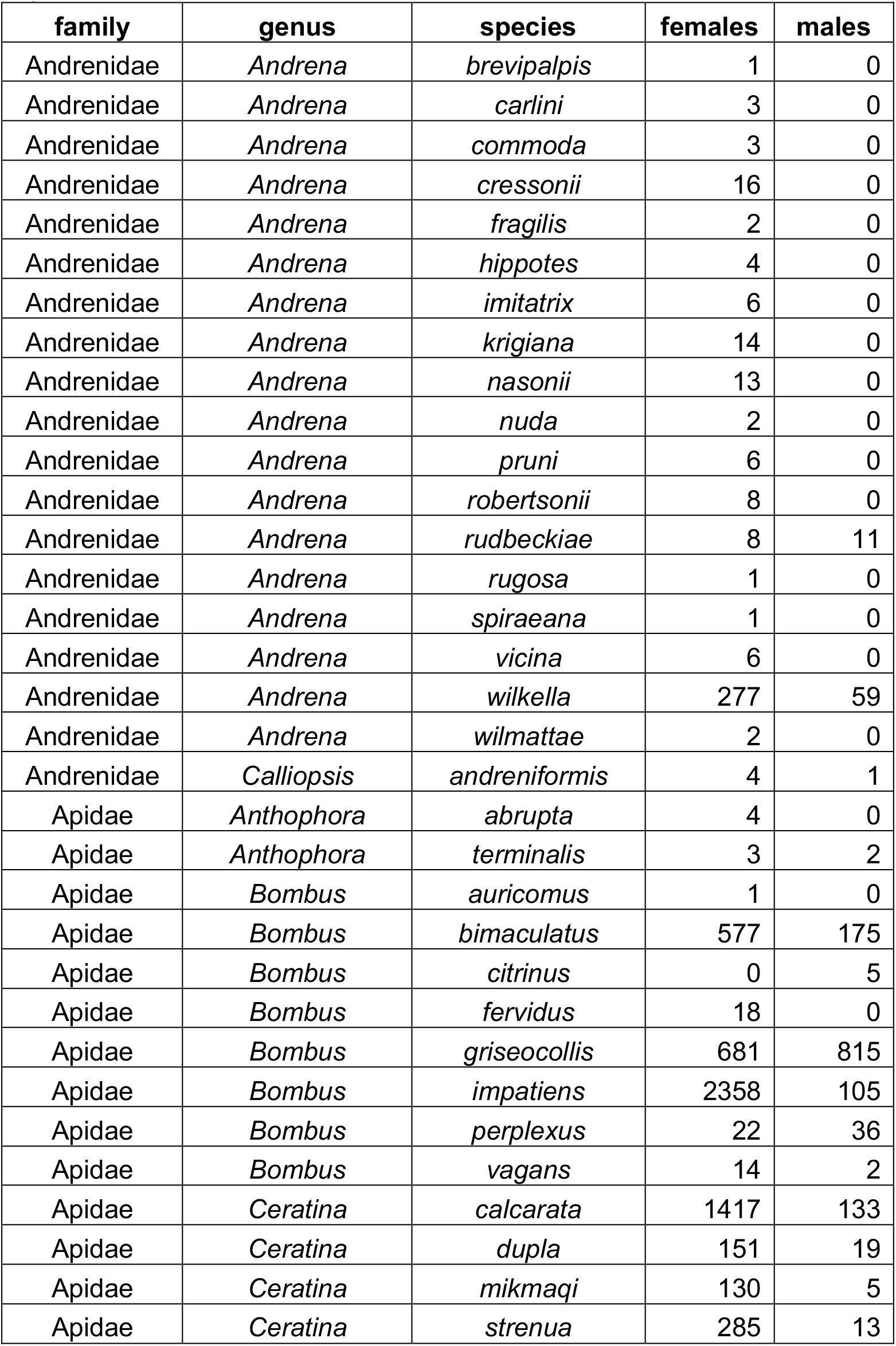
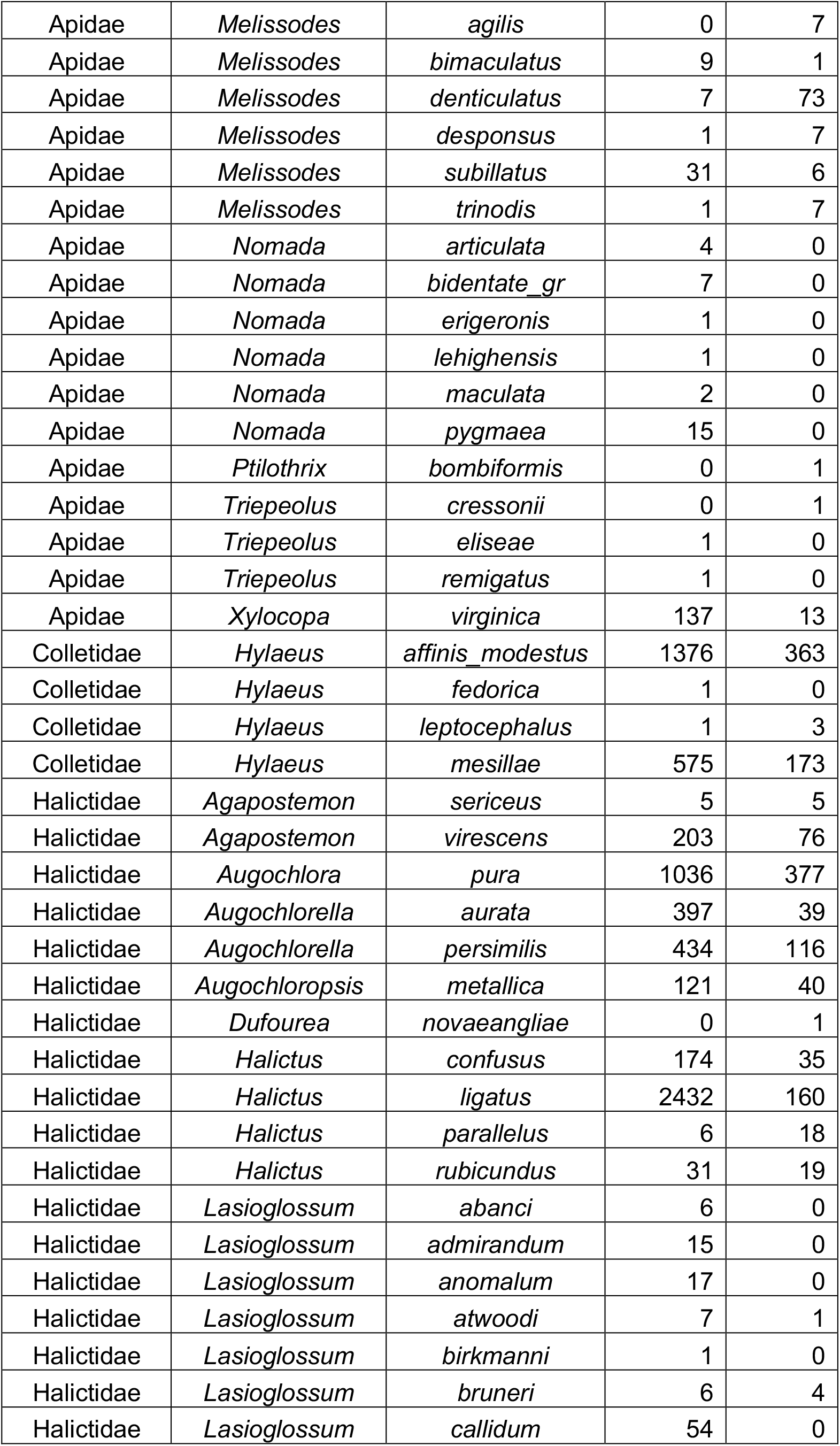
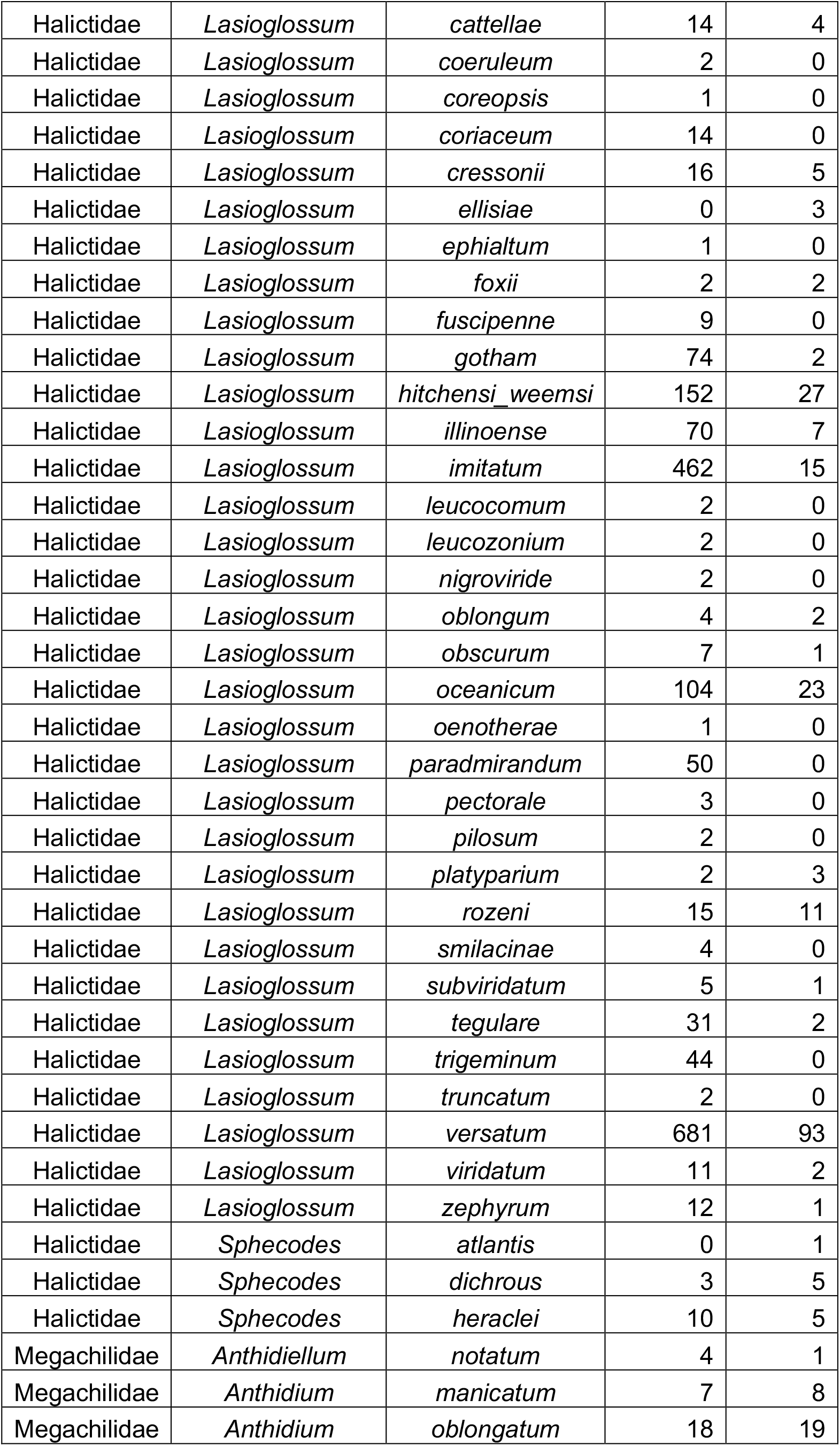
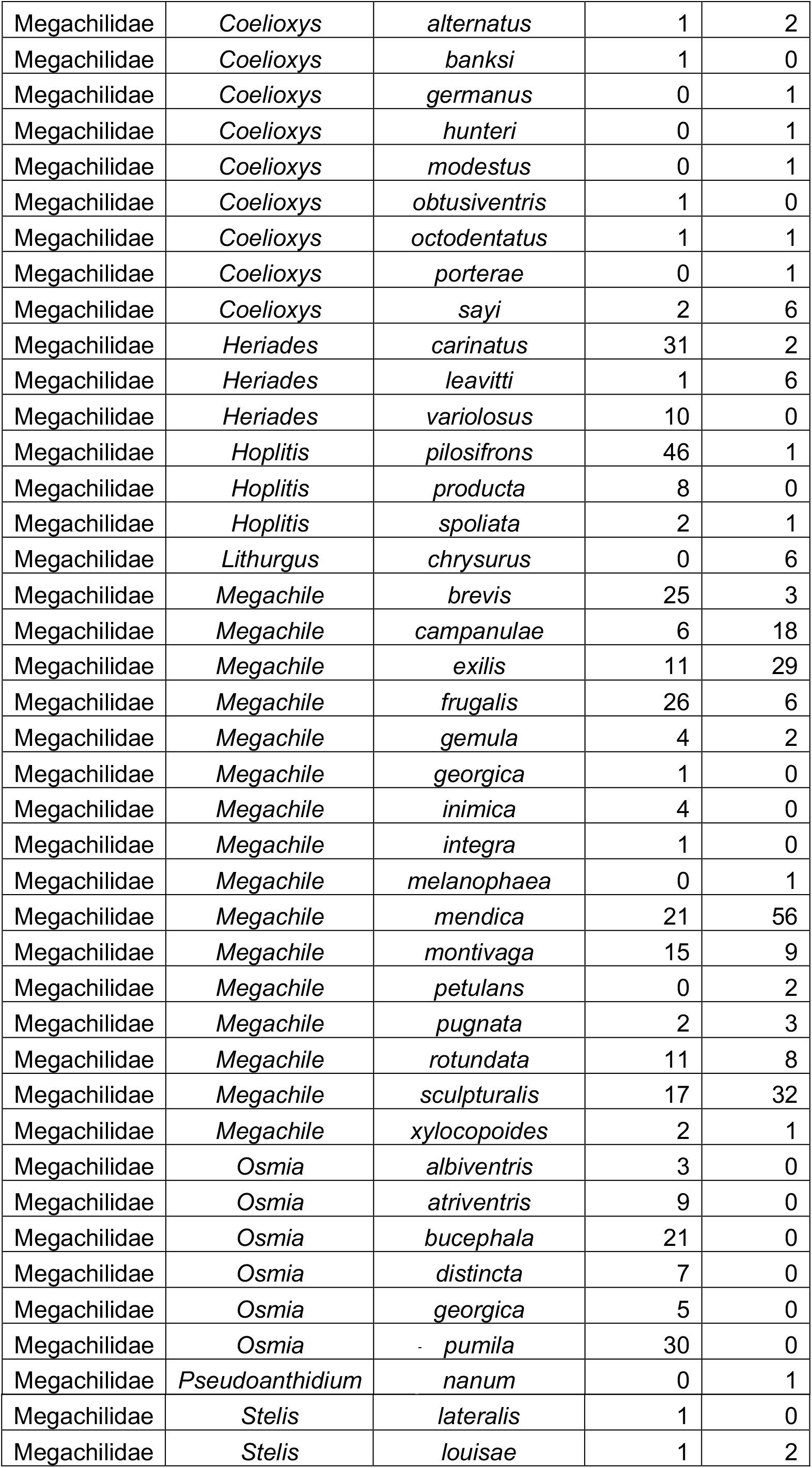
Bee species with number of female and male specimens collected. *This table will be removed from the final submission when data are deposited on Dryad*.

**Table S3.**
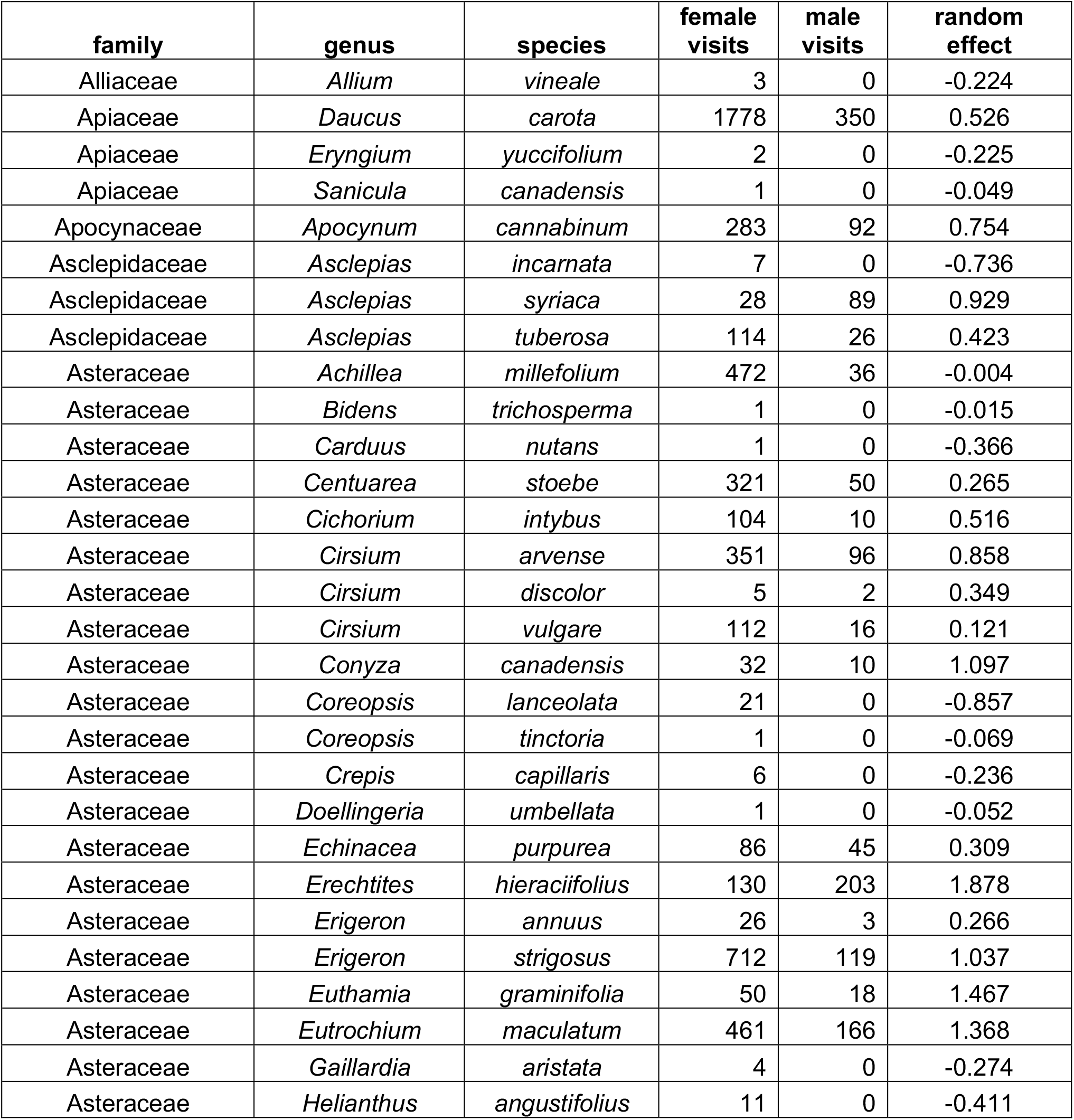
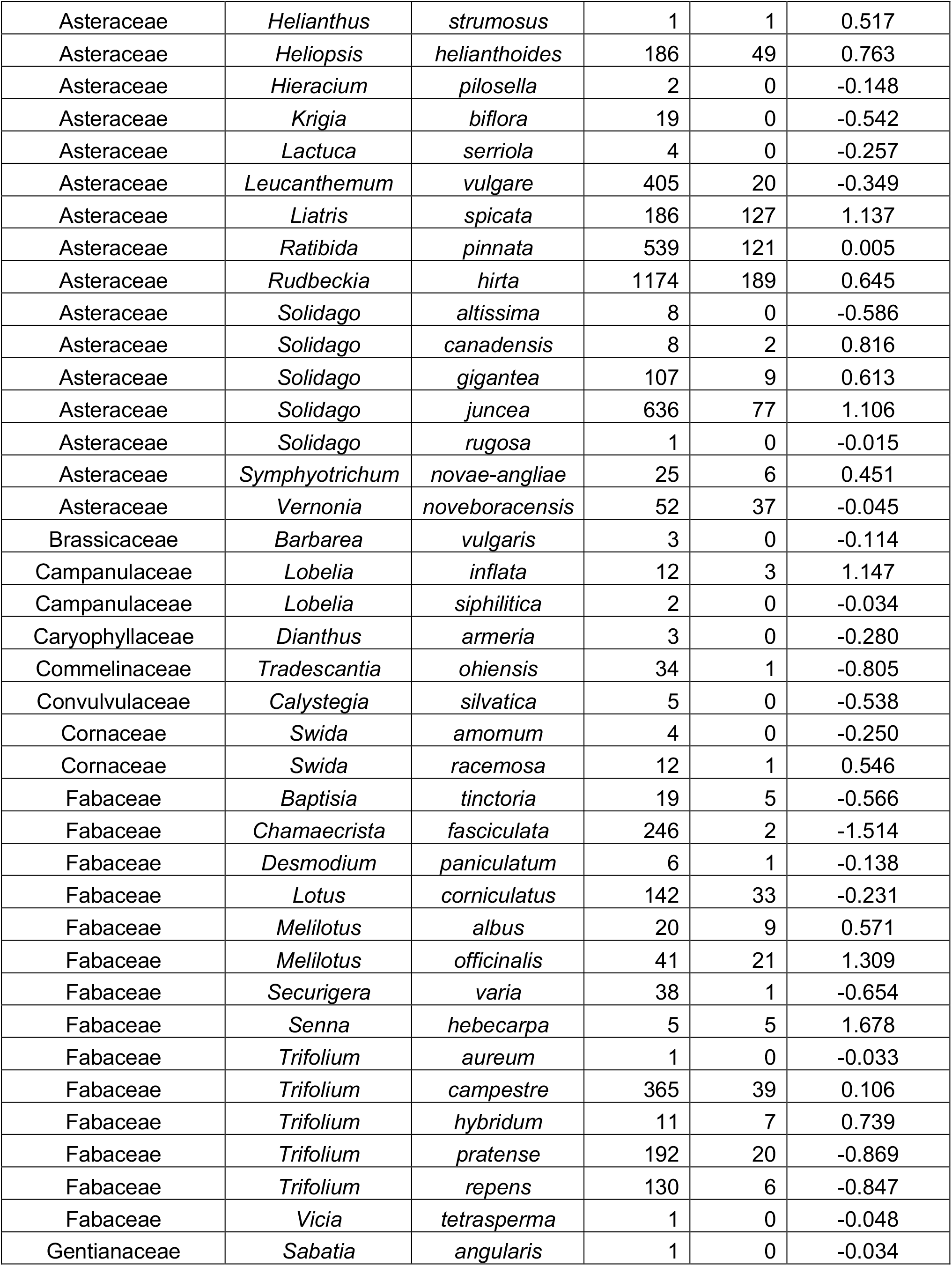
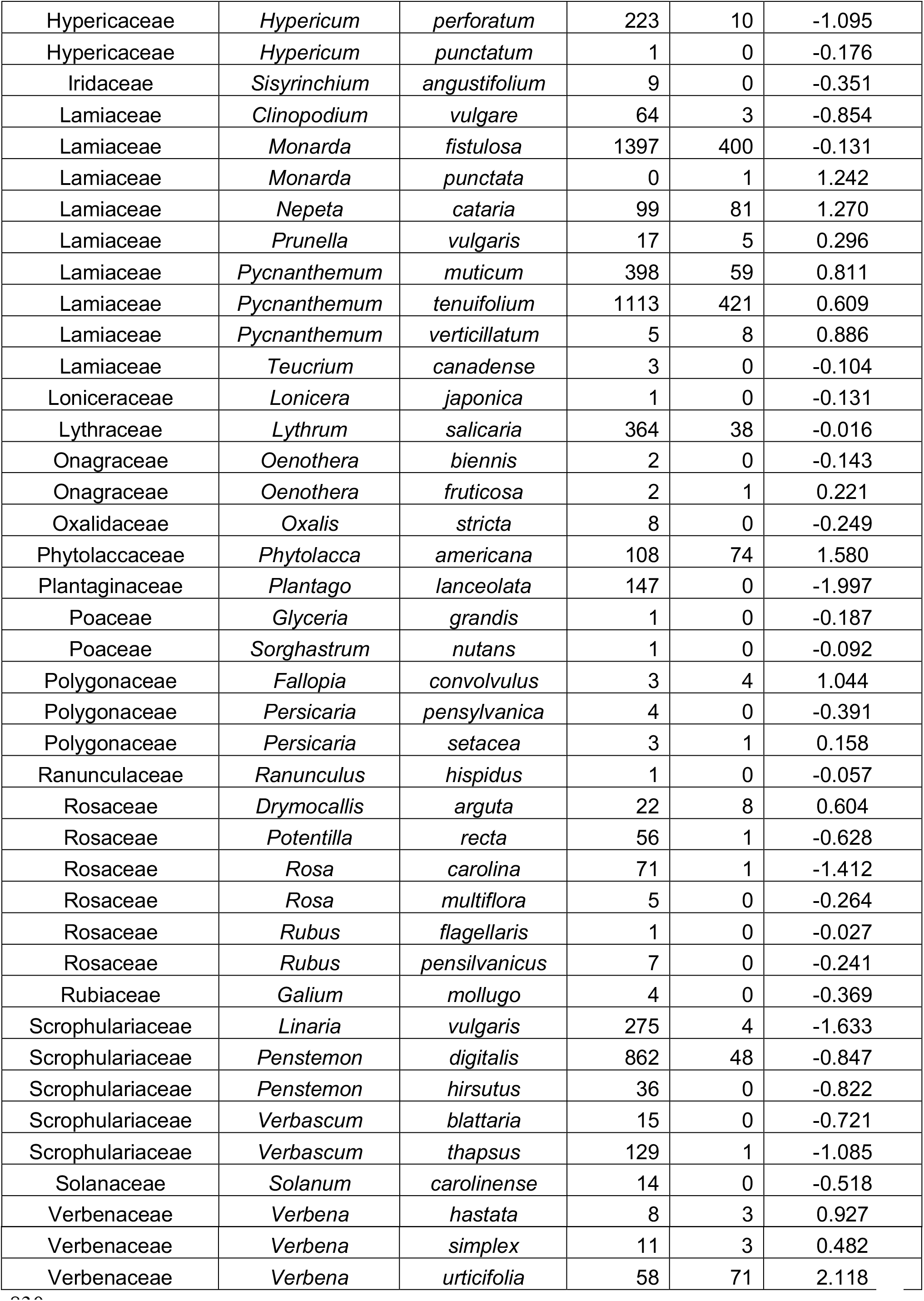
Number of male and female visitors to each plant species, and bias towards attracting male bee visitors. This bias is the random effect prediction from the seasonal model, which indicates the change in log(odds) that a visiting bee is male when the species of flower it visits is given; greater values indicate male bias.

